# Integrin-TGFβ axis induces partial EMT in basal-like cells to lead collective invasion

**DOI:** 10.1101/2025.04.04.647177

**Authors:** Abdelrahman G. El-Gammal, Lisa Jansen, Mathijs P. Verhagen, Hilverd Kremer, Sarmad Soufi, Mina Masoudnia, Isabella da Silva, Denise Westland, Peter Bult, Riccardo Fodde, Monique M. A. Verstegen, Martijn Gloerich, Saskia W. C. van Mil, Antoine A. Khalil

## Abstract

Collective invasion is the predominant mode of cancer cell dissemination in breast cancer and represents the initial step of metastatic spread. Basal-like leader cells drive this process by maintaining cell-cell junctions with the follower cells while extending actin-rich protrusions and remodeling the collagen I-rich peritumoral stroma. These features resemble those of individually-invading cells following epithelial-to-mesenchymal transition (EMT). However, how leader cells acquire these traits while preserving cohesion within the collective remains unclear. Here, we identify a collagen I-responsive subset of basal-like cells that coexpress cytoskeletal, extracellular matrix (ECM)-remodeling, and epithelial junction genes. We show that integrin α2 (Itgα2) links collagen I engagement to mesenchymal reprogramming by inducing inhibin beta A (INHBA) expression and activating tumor growth factor β (TGFβ) signaling. This, in turn, upregulates vimentin while preserving epithelial junction gene expression. In parallel, Itgα2 promotes ECM degradation through a TGFβ-independent mechanism. This study identifies Itgα2-TGFβ axis as a key regulator of partial EMT and leader cell function, highlighting it as a potential therapeutic target in aggressive breast cancers.

## Introduction

Invasive progression in epithelial tumors involves not only changes in cell motility, but also the emergence of coordinated multicellular behaviors. In breast cancer, tumor cells often disseminate collectively, invading as multicellular strands that maintain epithelial connectivity while navigating through the extracellular matrix (ECM) stroma^1,2^. This process is guided by leader cells, specialized cells at the invasive front that extend actin-rich protrusions into the ECM while remaining coupled to follower cells via stable junctions^3,4^. These leader cells exhibit features commonly associated with epithelial-to-mesenchymal transition (EMT), including cytoskeletal remodeling and matrix interaction, yet they retain epithelial architecture and mechanical integration within the collective^5–7^. EMT is a transcriptional program that enables epithelial cells to downregulate cell–cell adhesion, acquire mesenchymal traits, and become motile and invasive^4,8–10^. Although EMT has been extensively studied in the context of single-cell migration, how partial EMT arises in collective contexts remains poorly understood.

Leader cells represent a functional state within a subset of cells located at the tumor–stroma interface^3^. This state is not fixed but arises through dynamic regulation by local extracellular cues. In breast tumors, collagen I-rich ECM selectively promotes leader behavior in keratin-14–positive (K14⁺) basal-like cells^11,12^. These cells, enriched at the invasive front, drive collective invasion as multicellular strands. However, only a subset of basal-like cells acquires invasive functions, indicating that leader cell emergence is triggered by specific microenvironmental inputs. Given that basal-like cells not only mediate local invasion but also contribute to metastatic dissemination^13–15^, it is essential to define the molecular identity of the basal-like leader subset and to determine how ECM components such as collagen I instruct their reprogramming into an invasive leader state.

As tumors progress, the ECM undergoes substantial remodeling, including increased fibrillar collagen I deposition and stiffening, both of which correlate with poor prognosis in breast cancer patients^16–19^. The biochemical and physical changes in the ECM are sensed by cells through ECM receptors. Integrins, a family of transmembrane receptors composed of α and β subunits, are key mediators of ECM sensing^20–22^. Through their extracellular domain, integrins bind to ECM components, while their intracellular domains connect to the cytoskeleton via focal adhesion protein complexes^23^. Actomyosin-generated forces are transmitted through integrins to the ECM. In turn, the ECM exerts counteracting forces dictated by its stiffness and organization^24^. This dynamic force exchange is central to mechanotransduction, the process by which cells sense and respond to ECM rigidity and tension^8,23–25^. A key effector of this process is Yes-associated protein (YAP), which becomes activated in response to ECM stiffening^26^. We have previously shown that YAP is specifically induced in basal-like cells upon interaction with collagen I, where it drives a transcriptional program that promotes protrusive behavior and collagen fiber pulling and realignment^7^. This bidirectional interaction between basal-like cells and collagen I establishes mechanical and transcriptional feedback loops that reinforce leader cell emergence and invasion. However, whether interactions with collagen I also initiate mesenchymal transcriptional programs, and whether these traits are functionally required for leader cell behavior, remains unknown.

Here, we test whether collagen I interactions induce mesenchymal characteristics in basal-like cells to promote their transition into leader cells. Single-cell RNA-sequencing of breast cancer organoids identified a collagen-specific basal-like subpopulation enriched for EMT genes, including vimentin. Vimentin was also expressed in leader cells at the invasive front in invasive carcinoma of no special type (NST) patient tissues and was required for collective invasion in vitro. Mechanistically, we identified integrin α2 (Itgα2), part of the α2β1 receptor for fibrillar collagen I, as a key upstream regulator. Itgα2 controlled vimentin expression and ECM degradation via transforming growth factor β (TGFβ)-dependent and -independent pathways, respectively. Components of this pathway were associated with poor prognosis of breast cancer patients. These findings establish an integrin-based mechanism linking ECM interactions to mesenchymal reprogramming and collective invasion.

## Results

### A subset of basal-like cells with mesenchymal signature defines the leaders of collective invasion

To identify mesenchymal traits acquired by leader cells, we expanded our previous single-cell RNA-sequencing analysis on mouse mammary tumor virus (MMTV)-PyMT breast cancer organoids^27^. This model recapitulates key aspects of invasive carcinoma NST, including basal-guided collective invasion and lung metastasis^12,28^. When cultured in type I collagen, basal-like cells in these organoids acquire leader cell behavior and drive invasion as multicellular strands (Supplementary movie 1), whereas in basement membrane extract (BME), they remain non-invasive^11,12,27^. While our prior clustering distinguished basal- and luminal-like cell states, it did not resolve functionally distinct subsets within the basal compartment^27^.

Higher-resolution clustering revealed five distinct transcriptional clusters, including a breakdown of basal cells into two transcriptionally distinct subpopulations: a basal 1 cluster present in both BME and collagen I conditions, and a basal 2 cluster predominantly found in collagen I (Figure 1A, B). We also identified luminal-like populations, both cycling and non-cycling, and a distinct cluster enriched for G protein-coupled receptor kinase 4 (*Grk4*) and midline 1 (*Mid1*). Gene ontology analysis identified EMT as the top enriched gene set in basal 2 cells compared to basal 1 cells and the other populations (Figure 1C, S1A). In addition to EMT, basal 2 cells showed enrichment for hypoxia, inflammatory response, and tumor necrosis factor-alfa–Nuclear factor-κB (TNFα–NFκB) signaling pathways, all previously associated with invasive front states in breast cancer^29,30^. In contrast, basal 1 cells were enriched for oxidative phosphorylation and MYC target genes (Figure 1C) indicative of a metabolically active and transcriptionally proliferative phenotype^31,32^.

**Figure 1.**
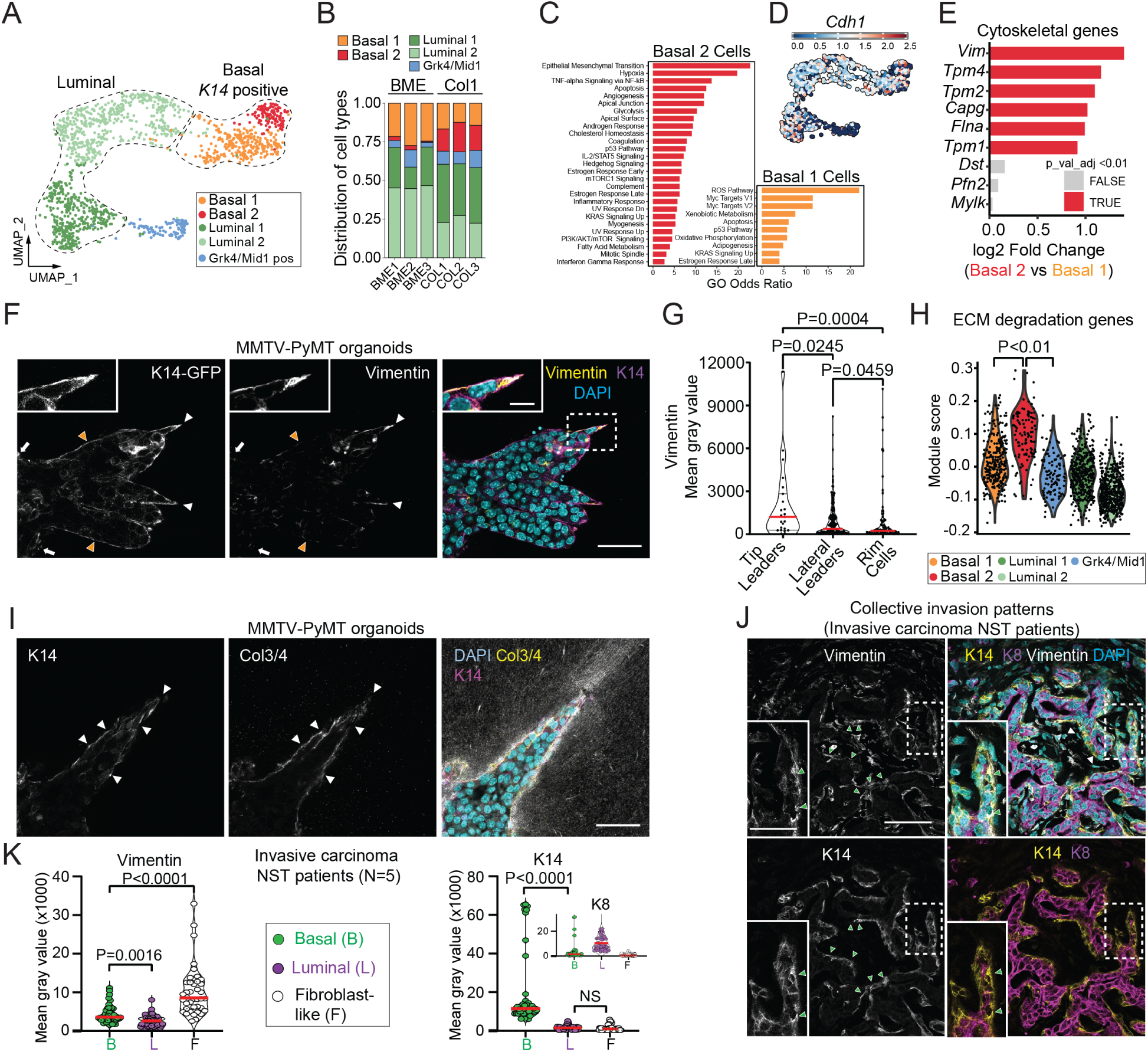
Leader cells emerge from a subset of basal-like cells possessing distinct mesenchymal features. **(A)** Uniform manifold approximation and projection (UMAP) plots based on transcriptional profiling from single-cell mRNA sequencing of MMTV-PyMT organoids cultured in 3D BME or collagen I (Khalil A. et al., 2024)27. (A) Distinct epithelial subpopulations were identified, including two K14-positive clusters (basal 1 and basal 2), Grk4/Mid1-positive cells, luminal 1, and luminal cycling (luminal 2). (**B**) Distribution of these epithelial subpopulations in organoids cultured in 3D BME (799 cells from 3 × 386-well plates) or collagen I (627 cells from 3 × 386-well plates). (**C**) Gene set enrichment analysis (GSEA) of basal 1 and basal 2 clusters showing significantly enriched pathways (filtered for NES > 0.5 and P < 0.01). (D) UMAP plot of Cdh1 expression across the epithelial subpopulations. (**E**) Log₂ fold changes in gene expression between basal 2 and basal 1 subsets focusing on genes belonging to the hallmarks of cancer gene set that encode cytoskeletal components, based on scRNA-seq data from MMTV-PyMT organoids. (**F**) Confocal imaging of K14-GFP and vimentin in MMTV-PyMT organoids in 3D collagen I, arrowheads depict vimentin-rich leader cells. (**G**) Mean-gray values of vimentin in K14-GFP+ cells located in different sites of the MMTV-PyMT organoids; tip leader cells (n = 23, white arrowheads), lateral leader cells (n = 179, orange arrowheads), and rim cells (n = 87, white arrows), from 3 independent experiments. (H) Module scores representing the aggregate expression of the ECM degradation gene set in individual cells. Scores reflect relative enrichment of ECM remodeling programs in basal 2 cells compared to the other subpopulations. (**I**) Immunostaining of K14 and col ¾ (degraded collagen I) in MMTV-PyMT organoids in 3D collagen I (reflection), arrowheads depict K14+ leader cells. (**J**) Confocal imaging (maximum projection) of K14, K8, and vimentin from a whole tissue section of a K14-positive invasive carcinoma NST patient (representative image of N=5 patients) with zoom in on protrusive leader cells expressing K14 and vimentin. Arrowheads: green: basal-like cells; white: fibroblast-like cells. (K) Mean-gray values of vimentin, K14, and K8 in the different cell types; basal-like (green, 45 cells), luminal-like (magenta, 44 cells) and fibroblast-like (white, 40 cells) from 5 invasive carcinoma NST patients. Zoom in depicting the K8 signal in the 3 populations. Scale bars: 50 µm (**F, I**), 10 µm (F, zoom-in), 100 µm (J), 25 µm (J, zoom-in). P values, two-sided Kruskal-Wallis test (Dunn’s multiple comparison) (G, K), one-way ANOVA with Tukey test for multiple comparisons (H).

Within the EMT gene set upregulated in the basal 2 subset were genes encoding for ECM components (*Fn1, Tnc, Ecm1, Lama3, Lamc2*), cell–ECM receptors (*Itgβ1, Sdc4*), cytoskeletal regulators (*Vim, Tpm1, Tpm2, Tpm4, Capg, Flna*), and TGFβ/activin-related signaling molecules (*Inhba, Tgfbi*) (Figure S1B). This mesenchymal signature was accompanied by preservation of epithelial junction genes, with most expressed at levels comparable to basal 1 and luminal cells, including *Cdh1* (E-cadherin), which is typically downregulated in full EMT (Figure 1D, S1C). Notably, basal 2 cells upregulated *Epcam* and downregulated *Tjp3*, but the overall junctional transcriptional profile remained intact, consistent with a partial EMT state.

Given the role of cytoskeletal remodeling in leader cell function, we examined cytoskeletal genes within the EMT gene set. *Vimentin* was among the most highly expressed genes across the EMT signature and was the most strongly upregulated cytoskeletal gene in basal 2 compared to basal 1 cells. In 3D collagen I, basal-like cells (K14⁺, K8-low) are located at the rim of the organoid (white arrows), at the tip (white arrowheads), and along the lateral edges (yellow arrowheads) of invasive strands (Figure S1D). Immunostaining for vimentin revealed that, within the basal-like cells, expression was highest in tip-positioned leader cells, followed by lateral strand leaders, and lowest in rim cells (Figure 1F, G). To test whether this observation extends beyond the PyMT model, we analyzed *Apc*^1572T^ organoids, which harbor an Apc mutation^33,34^ and are molecularly distinct from PyMT-driven tumors, but similarly recapitulate basal cell–guided (K14⁺) collective invasion when cultured in collagen I (Figure S1E). Similar to MMTV-PyMT, vimentin was consistently elevated in K14⁺ cells at the invasive front (Figure S1F). These findings strongly suggest that basal 2 cells correspond to the basal-like leader cells driving collective invasion of strands.

Because matrix degradation is a hallmark of mesenchymal single-cell invasion^35^, we next assessed whether basal 2 cells exhibit matrix-degrading potential. We analyzed scRNA-seq data using a selected gene set of ECM-degrading enzymes and their regulators (Supplementary information 1). While no single gene was dominantly enriched, *Mmp3*, *Mmp13*, and *Mmp14* showed a trend of higher expression in basal 2 cells (Figure S1G). This dispersed expression pattern reflects the known redundancy among ECM proteases, where degradation typically results from the combined activity of multiple enzymes^36^. Gene set enrichment analysis confirmed that ECM-degradation signatures were significantly enriched in basal 2 compared to basal 1 and non-basal populations (Figure 1H), indicating that matrix remodeling is a transcriptional feature of this collagen I-induced subset. To functionally validate this transcriptional signature, we performed immunostaining for the collagen ¾ (Col¾) epitope, a fragment generated by MMP-mediated cleavage of collagen I^37–39^. Confocal microscopy of MMTV-PyMT organoids cultured in collagen I for 3 days revealed strong Col¾ accumulation at the tumor-ECM interface, including the fronts and lateral edges of K14⁺ strands (Figure 1I, arrowheads). Although this signal does not pinpoint the source of proteolytic activity, its accumulation confirms active collagen I remodeling at sites of basal cell–ECM contact. The degrading abilities, together with the pulling abilities (Supplementary movie 1), support the matrix-remodeling potential of leader cells and are consistent with previous reports of ECM remodeling during mesenchymal single-cell invasion^35^.

To validate the mesenchymal character of leader cells in human tumors, we analyzed invasive carcinoma NST patient sections containing K14⁺ invasive fronts (N = 5). As in organoids, basal-like cells (K14-high, K8-low) (Figure 1J, green arrowheads) surrounded luminal-like cells (K8-high, K14-low) at the tumor-stroma interface (Figure 1J, S1H). Vimentin was strongly expressed in basal-like cells compared to adjacent luminal cells, though lower than in surrounding stromal cells (Figure 1K).

These data identify a basal-like subset with mesenchymal features and retained epithelial junctions, consistent with a hybrid epithelial–mesenchymal state that corresponds to leader cells driving collective invasion.

### Vimentin and ECM degradation are required for leader cell emergence and are regulated by the collagen I receptor Itgα2β1

Vimentin has recently been shown to be important for metastatic progression in breast cancer, but its specific role in driving invasion, particularly leader cell function, remains unclear^40^. To test whether vimentin is necessary for basal-like cells to develop into leader cells, we generated doxycycline (Dox)-inducible shRNA knockdown organoids targeting vimentin. Dox treatment reduced vimentin expression by >75% (shRNA1) and >65% (shRNA2) in MMTV-PyMT organoids (Figure S2A), and by >95% in *Apc*^1572T^ organoids (Figure S2B). Vimentin knockdown led to a marked reduction in invasion, as indicated by decreased leader cell numbers per organoid (Figure 2A, B; S2C–E) and shorter invasive strands (Figure S2F, G). In MMTV-PyMT organoids, vimentin knockdown also reduced organoid body size (Figure S2H), an effect not observed in *Apc*^1572T^ organoids (Figure S2I), suggesting a model-specific role in tumor growth. Together, these findings demonstrate that vimentin is essential for the emergence of basal-like leader cells and for the extension of invasive strands, establishing it as a critical mesenchymal effector of collective invasion.

**Figure 2.**
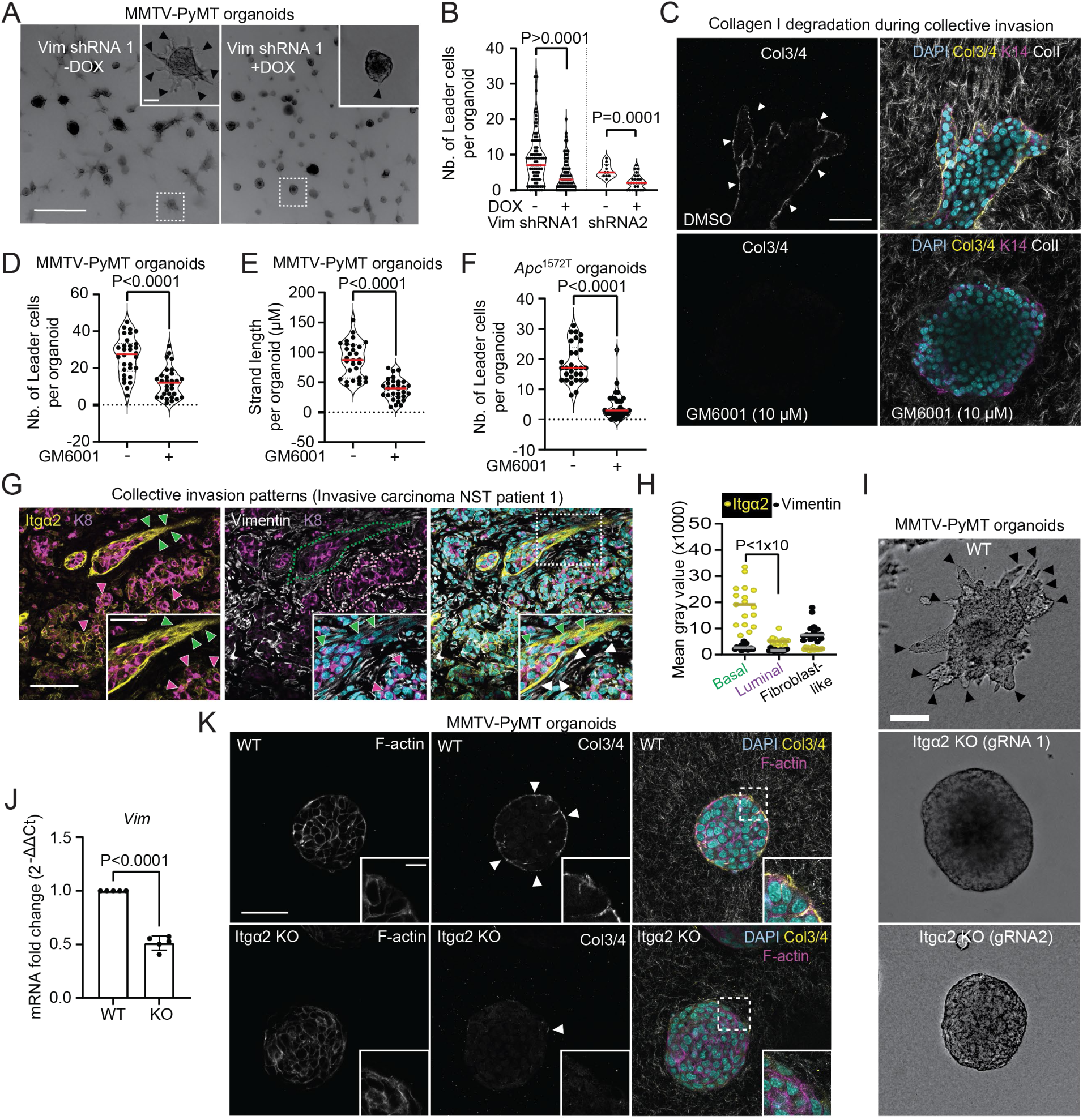
Leader cell function and collective invasion rely on vimentin expression and ECM degradation. **(A)** Vimentin shRNA (−/+ Dox) in MMTV-PyMT organoids grown in collagen I for 3 days. Arrowheads: protrusive leader cells. **(B)** Quantification of number of leader cells per organoid in MMTV-PyMT: *n* = 87 (– DOX), *n* = 87 (+DOX) for shRNA1, and *n* = 29 (−DOX), *n* = 44 (+DOX) for shRNA2, from three (shRNA1) and two (shRNA2) independent experiments. **(C)** Confocal imaging of K14 and col ¾ (degraded collagen I) in MMTV-PyMT organoids treated with GM6001 or DMSO control for 3 days in collagen I. Arrowheads: pericellular signal of col ¾ adjacent to basal-like cells (K14). **(D, E)** Quantification of **(D)** number of leader cells and **(E)** strand length per organoid in MMTV-PyMT organoids. Red line indicates the median, from *n* = 30 organoids per condition from three independent experiments. **(F)** Number of leader cells per organoid in *Apc*^1572T^ organoids cultured in 3D collagen I for 3 days and treated with GM6001 or DMSO control. Red line indicates the median, from *n* = 30 organoids per condition from three independent experiments. **(G)** Confocal imaging of Itgα2, vimentin, and K8 in an invasive carcinoma NST patient tissue section. Green contour: cluster of cancer cells with protrusive morphology; green arrowheads: basal-like cells (K8-low) at the tumor–stroma interface with high Itgα2 expression. White contour: cluster of cancer cells lacking basal-like cells (K8-high), with low Itgα2 expression; magenta arrowheads. White arrowheads: fibroblast-like cells (elongated, spindle-shaped). **(H)** Quantification of mean gray values for Itgα2 and vimentin in basal-like (*n* = 33), luminal-like (*n* = 32), and fibroblast-like (*n* = 32) cells from one invasive carcinoma NST patient tissue section. **(I)** Representative brightfield images of MMTV-PyMT organoids (Itgα2-WT or Itgα2-KO, gRNA1 and gRNA2) cultured in 3D collagen I. Black arrowheads: invasive strands. **(J)** qPCR analysis of *Vim* mRNA expression in Itgα2-KO relative to Itgα2-WT in MMTV-PyMT organoids cultured in 3D collagen I for three days. Values represent mean normalized mRNA expression (relative to housekeeping genes). Data are presented as mean ± SD from five independent experiments. **(K)** Confocal imaging of col ¾ and F-actin in Itgα2-WT and Itgα2-KO MMTV-PyMT organoids after one day in 3D collagen I. Scale bars: 1000 μm **(A)**; 10 μm (**A**, zoom-in); 50 μm **(C, K)**; 10 μm (**K**, zoom-in); 100 μm **(G, I)**; 50 μm (**G**, zoom-in). P values: two-sided unpaired Mann–Whitney test **(B, D, E, F)**; two-sided Student’s *t* test **(J)**; two-sided Kruskal-Wallis test with Dunn’s multiple comparisons **(H)**.

We also assessed the role of ECM degradation in collective invasion. Treatment with GM6001, a broad-spectrum matrix metalloproteinase (MMP) inhibitor, markedly reduced Col¾ signal compared to controls (Figure 2C). MMP inhibition significantly impaired collective invasion in both MMTV-PyMT and *Apc*^1572T^ organoids, reducing leader cell numbers and strand elongation by more than 50% (Figure 2D-F; S2J, K), without affecting organoid body size (Figure S2L, M). These findings show that collagen I degradation is required for both leader cell emergence and sustained collective invasion in 3D collagen I.

Since mesenchymal characteristics are promoted in collagen I environments, we next asked whether these features are driven by direct interaction with the ECM. We focused on Itgα2, the collagen-binding subunit of the Itgα2β1 heterodimer, which has the highest known affinity for fibrillar collagen I^41–45^. *Itgβ1* mRNA was upregulated in basal 2 cells compared to the other basal and luminal cells. While *Itgα2* mRNA showed only a modest trend toward higher expression in basal 2 cells (Figure S2N), it was strongly expressed at the protein levels in basal-like cells (K8-low) at the tips of invasive strands compared to the signal observed in the follower cells (membranous/cell to cell contact) (Figure S2O). This pattern was confirmed in invasive carcinoma NST patient samples, where Itgα2 was enriched in basal-like cells (K8-low) at the tumor-stroma interface, in contrast to lower expression in K8-high luminal cells and fibroblast-like cells (Figure 2G, H). Together, these data indicate that Itgα2 is expressed in leader-positioned basal-like cells and likely mediates tumor-collagen I interactions during collective invasion.

Because Itgβ1 partners with multiple α-integrins and its deletion broadly affects cell proliferation and survival across cell types^46,47^, we chose to focus on Itgα2 to specifically assess collagen I-dependent functions. CRISPR/Cas9-mediated deletion of *Itgα2* in MMTV-PyMT organoids (Figure S2P, Q) abolished collective invasion in collagen I (Figure 2I; S2R), preventing basal-like cells from protruding and forming invasive strands (Figure S2O, white arrows). Loss of Itgα2 also markedly reduced vimentin expression (Figure 2J) and impaired collagen degradation (Figure 2K; S2S), demonstrating that collagen I-Itgα2β1 engagement is required to activate mesenchymal traits in basal-like cells and to drive collective invasion.

### TGFβ signaling promotes vimentin expression and collective invasion

TGFβ signaling is a major inducer of EMT that drives single cell invasion^48–50^. Given the mesenchymal signature of basal-like leader cells, we next investigated whether TGFβ signaling contributes to this collagen I-induced gene expression program. TGFβ signaling is initiated by ligand binding to type II receptors, which activate type I receptors, including ALK4, ALK5, and ALK7, thereby triggering downstream SMAD-dependent transcriptional responses^51,52^. Gene set enrichment analysis (KEGG) indicated elevated TGFβ pathway activity in the collagen I-specific basal 2 population (Figure S3A). We further analyzed the expression of a selected set of canonical TGFβ target genes (Supplementary information 1), including *Ctgf*, *Smad7* and *Pai1*, and found this gene set to be also strongly upregulated in basal 2 cells compared to basal 1 subset and the other populations (Figure 3A), suggesting active TGFβ signaling in leader cells in collagen I. To test whether TGFβ regulates mesenchymal gene expression, we treated organoids cultured in collagen I with SB431542 (5 µM), a selective inhibitor of TGFβ type I receptor kinases (ALK4, ALK5 and ALK7)^53,54^ and analyzed gene expression by qPCR (Figure 3B). SB431542 treatment significantly reduced the expression of canonical TGFβ target genes, including *Ctgf* and *Smad7*, confirming effective pathway inhibition. In MMTV-PyMT organoids, expression of the EMT effector *Vim* and the transcriptional regulator *Slug* was also reduced, while *Snai1* mRNA levels remained unchanged. Similarly, SB431542 treatment of *Apc*^1572T^ organoids reduced *Vim* and *Ctgf*, while changes in *Snai1* and *Smad7* were not significant (Figure S3B). Interestingly, TGFβ inhibition did not impair ECM degradation. Immunofluorescence for the collagen ¾ (Col¾) epitope showed comparable levels in SB431542-treated and control organoids (Figure 3C). These findings were corroborated using A83-01 (0.5 µM), a more potent ALK inhibitor (Figure S3C, D) indicating that matrix degradation is TGFβ-independent. These findings show that TGFβ signaling is transcriptionally active in collagen I-embedded basal-like cells and contribute to the mesenchymal gene profile.

**Figure 3.**
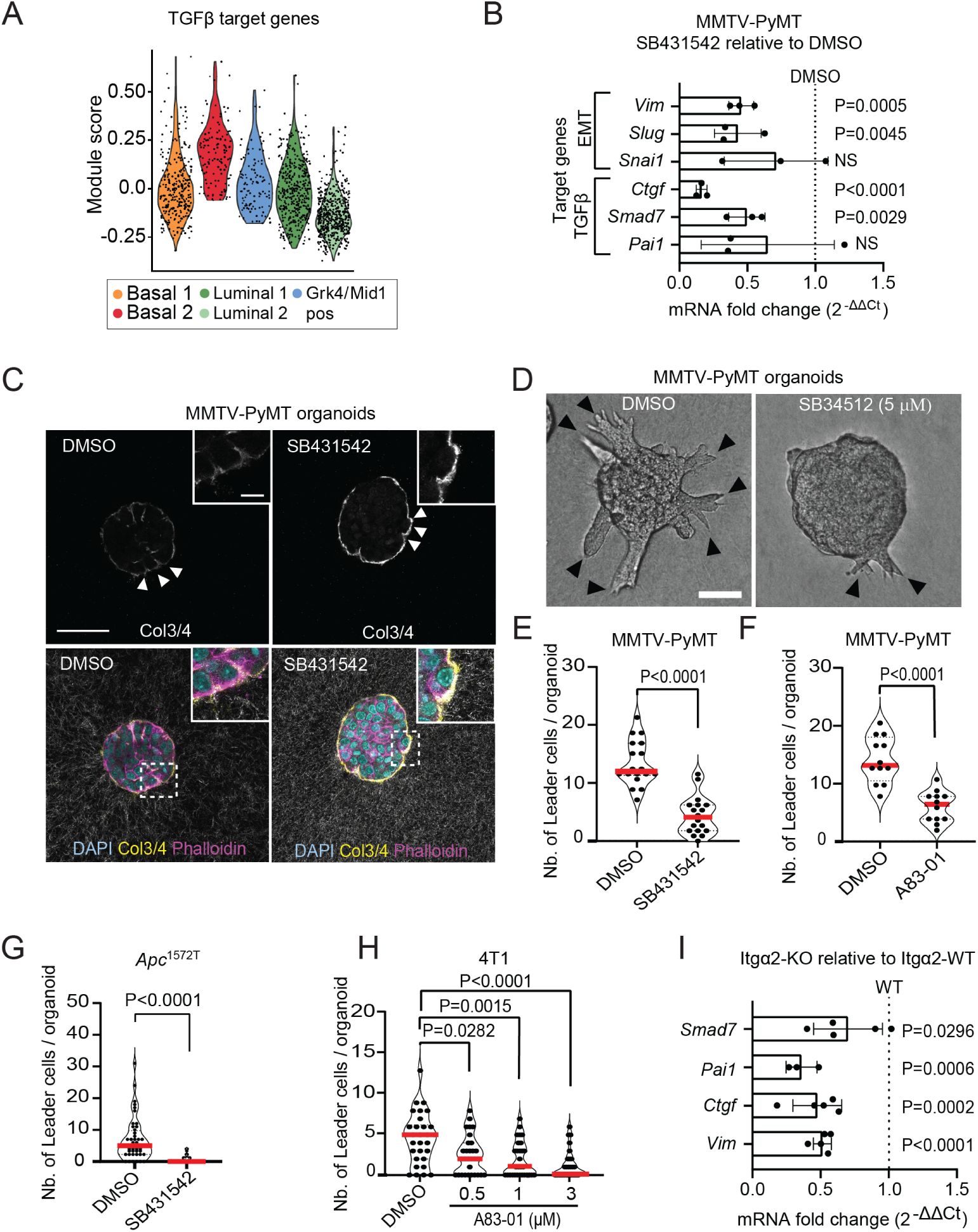
TGFβ signaling drives collective invasion and vimentin expression. **(A)** Module scores representing the aggregate expression of TGFβ target genes in individual cells, reflecting relative pathway enrichment in basal 2 cells compared to the other subpopulations. **(B)** qPCR analysis of classical TGFβ and EMT target genes in MMTV-PyMT organoids cultured in 3D collagen I for three days. Values represent mean normalized mRNA expression (relative to housekeeping genes) in organoids treated with 5 μM SB431542, shown relative to DMSO-treated controls (dashed line). Data represent mean ± SD from three independent experiments. **(C)** Confocal imaging of col ¾ (degraded collagen I) and F-actin in MMTV-PyMT organoids cultured in 3D collagen I and treated with SB431542 (5 μM) or DMSO control for one day. Arrowheads: pericellular col ¾ signal. Inset: organoid-collagen I interface. **(D)** Representative brightfield images of MMTV-PyMT organoids cultured in 3D collagen I and treated with SB431542 or DMSO control for three days. **(E-H)** Quantification of number of leader cells per organoid in **(E, F)** MMTV-PyMT organoids treated with **(E)** 5 μM SB431542 or **(F)** 0.5 μM A83-01 or DMSO control, **(G)** *Apc*^1572T^ organoids treated with 5 μM SB431542 or DMSO control, and **(H)** 4T1 cancer spheroids treated with a dose series of A8301 (0.5 μM, 1 μM, 3 μM). Red line indicates the median from **(E)** *n =* 18 organoids, **(F)** *n =* 12 organoids, **(G)** *n =* 44 organoids, **(H)** *n =* 30 spheroids from three independent experiments. **(I)** qPCR analysis of classical TGFβ and EMT target genes in Itgα2-WT and Itgα2-KO MMTV-PyMT organoids cultured in 3D collagen I for three days. Values represent mean normalized mRNA expression (relative to housekeeping genes), shown for KO organoids relative to WT controls (dashed line). Data are presented as mean ± SD from three independent experiments. Scale bars: 50 μm **(C)**; 10 μm (**C**, zoom-in); 100 μm **(D)**. P values: one-way ANOVA with Tukey’s test for multiple comparisons **(A)**; two-sided Student’s *t* test **(B,I)** ;two-sided unpaired Mann–Whitney test **(E, F, G)**; two-sided Kruskal-Wallis test with Dunn’s multiple comparisons **(H)**.

Functionally, both SB431542 and A83-01 significantly reduced leader cell formation in MMTV-PyMT (Figure 3D-F) and *Apc*^1572T^ (Figure 3G) organoids. To test whether this requirement for TGFβ signaling extends to other breast cancer models, we examined 4T1 spheroids, a metastatic murine cell line known to invade collectively in 3D collagen I^55^. TGFβ inhibition in 4T1 spheroids similarly impaired leader cell emergence (Figure 3H; S3E, F). While TGFβ inhibition did not affect the overall size of MMTV-PyMT organoids or 4T1 spheroids (Figure S3G-J), it led to a reduction in *Apc*^1572T^ organoid size compared to Dimethyl sulfoxide (DMSO)-treated controls (Figure S3K), suggesting that TGFβ may have broader roles in tumor growth in this model.

Given that TGFβ inhibition and *Itgα2* loss produced similar effects on leader cell emergence and mesenchymal gene expression, we tested whether these pathways are functionally linked. qPCR analysis of breast cancer organoids cultured in 3D collagen I revealed that Itgα2 knockout significantly reduced the expression of TGFβ target genes, including *Ctgf*, *Pai1*, and *Smad7* (Figure 3I), indicating that Itgα2 promotes TGFβ pathway.

Our results demonstrate that collagen I-integrin interactions promote TGFβ signaling which is essential for leader cell formation and vimentin expression but is dispensable for matrix degradation.

### Collagen I-Itgα2β1 interactions promote mesenchymal reprogramming and TGFβ signaling via Inhba

To identify the ligand responsible for activating TGFβ signaling in basal-like cells within collagen I environments, we analyzed the expression of a selected panel of TGFβ superfamily ligands, including canonical *Tgfb1–3*, *Inhba–e*, *Bmp2-15*, *Gdf1-15*, *Nodal*, *Lefty1/2*, *Amh*, and *Thbs1*. Among these, *Inhba* was the most strongly upregulated ligand gene in basal 2 cells compared to basal 1 (Figure 4A). *Inhba* encodes the βA subunit of the homodimer activin A, a functional TGFβ ligand that activates SMAD-dependent transcriptional programs^56,57^. Consistent with its transcriptional regulation by collagen I-Itgα2 signaling, *Inhba* was significantly downregulated in *Itgα2*-deficient organoids at both mRNA and protein levels (Figure 4B, C; S4A).

**Figure 4.**
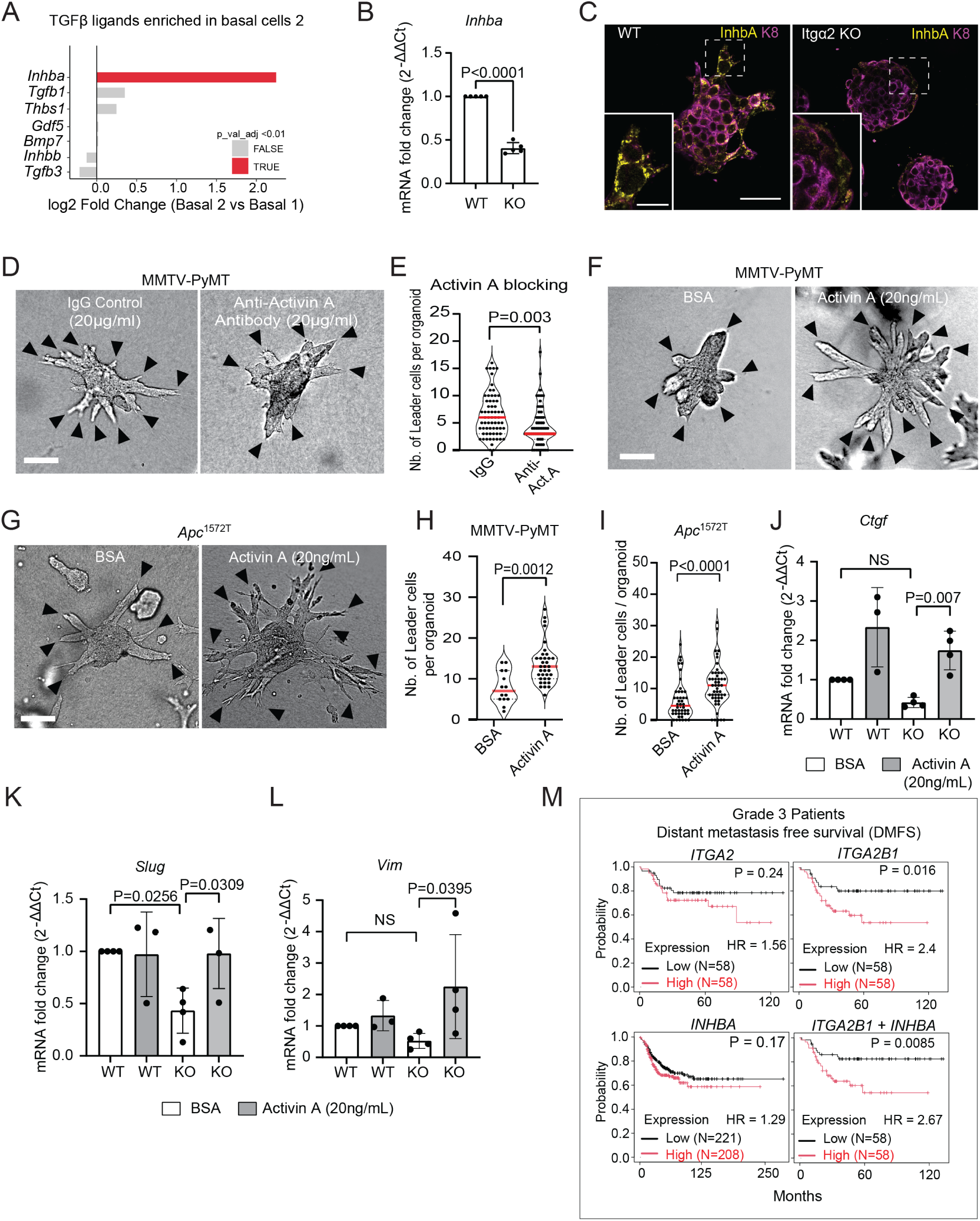
TGFβ signaling and mesenchymal traits are promoted by collagen I-Itgα2β1 interactions via Inhba. **(A)** Log₂ fold change of TGFβ ligands in the basal 2 versus basal 1 subset, based on scRNA-seq data from MMTV-PyMT organoids. The gene set includes ligands from all major TGFβ superfamily branches, including canonical TGFβs, activins/inhibins, BMPs, nodal-related ligands, and GDFs. **(B)** PCR analysis of *Inhba* mRNA expression in Itgα2-KO relative to Itgα2-WT in MMTV-PyMT organoids cultured in 3D collagen I for three days. Values represent mean normalized mRNA expression (relative to housekeeping genes). Data are presented as mean ± SD from five independent experiments. **(C)** Single confocal slice showing Inhba and K8 expression in Itgα2-WT and Itgα2-KO MMTV-PyMT organoids cultured in 3D collagen I for three days. Insets show basal-like cells (K8 low) guiding invasive strands (Itgα2-WT) or remaining at the organoid rim (Itgα2 KO, non-invasive). **(D)** Brightfield images of MMTV-PyMT organoids treated with an activin A neutralizing antibody (20 μg/ml) or IgG control (20 μg/ml) for two days in 3D collagen I. **(E)** Quantification of leader cell numbers per organoid under activin A neutralization or IgG control conditions. Red lines indicate the median; *n* = 60 organoids per condition from three independent experiments. **(F, G)** Brightfield images of **(F)** MMTV-PyMT and **(G)** *Apc*^1572T^ organoids treated with activin A ligand (20 ng/μl) or vehicle control (0.1% BSA) for three days in 3D collagen I. Arrowheads **(D, F, G)**: leader cells. **(H, I)** Quantification of leader cells per organoid in **(H)** MMTV-PyMT and **(I)** *Apc*^1572T^ organoids treated with activin A ligand (20 ng/μl) or vehicle control (0.1% BSA). Red line indicates the median from **(H)** *n* = 14–36 organoids per condition and **(I)** *n* = 44 organoids per condition from three independent experiments. **(J-L)** qPCR analysis of **(J)** *Ctgf*, **(K)** *Slug*, and **(L)** *Vim* mRNA expression in Itgα2-KO versus Itgα2-WT MMTV-PyMT organoids treated with activin A (20 ng/μl) or vehicle control (0.1% BSA) for three days. Bar graphs represent mean normalized expression values ± SD from three to four independent experiments. **(M)** Kaplan–Meier analysis correlating high vs. low mRNA expression of *INHBA*, *ITGA2*, *ITGA2B1*, and the combination of *ITGA2B1* + *INHBA* with distant metastasis-free survival (DMFS) in patients with grade 3 breast cancer. Scale bars: 50 μm **(C)**; 25 μm (**C**, Zoom-in); 100 μm **(D, F, G)**. P values: two-sided Student’s *t* test **(B)**; two-sided unpaired Mann–Whitney test **(E, H, I)**; one-way ANOVA with Šidák’s test for multiple comparisons **(J, K, L)**; Log-rank test **(M)**.

To test whether activin A contributes functionally to collective invasion, we neutralized endogenous activin A using a blocking antibody, which significantly reduced leader cell numbers per organoid compared to IgG-treated controls (Figure 4D, E). Conversely, supplementation with exogenous activin A (20 ng/ml) increased leader cell emergence in MMTV-PyMT organoids (1.75-fold), *Apc*^1572T^ organoids (2-fold), and 4T1 spheroids (1.8-fold) relative to BSA-treated controls (Figure 4F–I; S4B).

We next tested whether activin A could restore TGFβ signaling in the absence of Itgα2. Treatment of *Itgα2*-deficient organoids with exogenous activin A restored expression of the TGFβ target gene *Ctgf* but not *Smad7* (Figure 4J, S4D), and induced mesenchymal genes including *Vim*, even in the absence of Itgα2 (Figure 4K, L). However, activin A did not rescue collective invasion (Figure S4C).

Given that basal-like leader cells exhibit a partial EMT transcriptional profile, we next examined whether collagen I-Itgα2 signaling modulates *Cdh1* expression. Bulk qPCR of MMTV-PyMT organoids revealed a ∼1.7-fold increase in *Cdh1* mRNA upon *Itgα2* knockout (Figure S4E), consistent with a repressive input from collagen I-Itgα2 signaling. In contrast, single-cell RNA sequencing showed that *Cdh1* transcript levels in leader cells (in contact with collagen I), remained comparable to other cancer cell populations including luminal cells (Figure 1D, S1C). This suggests that *Itgα2*-dependent repression of E-cadherin is counteracted in leader cells, likely by additional transcriptional inputs active in the collagen-rich niche.

To test whether TGFβ signaling contributes to *Cdh1* regulation, we pharmacologically modulated the pathway. Inhibition with SB431542 suppressed canonical EMT targets but had no significant effect on *Cdh1* levels (Figure S4E). In contrast, activin A treatment led to a modest but reproducible increase in *Cdh1* mRNA (Figure S4E). This co-induction of mesenchymal programs alongside sustained *Cdh1* expression reflects the context-dependent role of activin A in EMT and is supported by prior studies showing that activin A signaling can maintain or even enhance E-cadherin expression in other epithelial cancer models^58,59^. Together, these findings indicate that collagen I-Itgα2 signaling promotes mesenchymal gene expression while sustaining *Cdh1*, at least in part, via activin A, supporting a transcriptional program consistent with partial EMT in basal-like leader cells.

To assess the clinical relevance of the Itgα2β1-Inhba signaling axis in breast cancer progression, we analyzed associations between gene expression and distant metastasis-free survival (DMFS) using the Györffy cohort of grade 3 breast cancer patients^60^. Individually, *ITGA2* and *ITGB1* expression levels showed no significant association with outcome (Figure 4M, S4F). However, their combined high expression, suggestive of increased integrin α2β1 receptor expression, was significantly associated with reduced metastasis-free survival (hazard ratio = 2.4, *P* = 0.016). Similarly, *INHBA* expression alone did not associate with DMFS, but when co-expressed with high *ITGA2B1*, the association with poor outcome was further strengthened (hazard ratio = 2.67, *P* = 0.0085) (Figure 4M). These data suggest that concurrent high expression of *ITGA2*, *ITGB1*, and *INHBA* may define a patient subgroup with markedly worse metastatic outcomes, highlighting the clinical relevance of the Itgα2β1-Inhba signaling axis.

Summarized, our findings demonstrate that collagen I engagement via Itgα2 induces upregulation of the TGFβ ligand Inhba, thereby activating the TGFβ pathway and promoting mesenchymal reprogramming in leader cells. This integrin-TGFβ cascade links ECM interactions to partial EMT and collective invasion, establishing it as a central driver of metastatic progression and a potential therapeutic target in aggressive breast cancer.

## Discussion

Our study identifies a collagen I-Itgα2-driven program through which leader cells acquire selective mesenchymal traits required for collective invasion. We show that Itgα2 links collagen I sensing to vimentin expression and matrix degradation, establishing a feedback loop that sustains leader cell function. This program enables basal-like cells to invade while maintaining epithelial cohesion, representing a hybrid epithelial–mesenchymal state distinct from full EMT. These findings reveal how ECM engagement and autocrine/paracrine signaling converge to generate a progressively evolving microenvironment that supports and drives collective invasion in aggressive breast cancers.

Using transcriptional profiling and functional assays in breast cancer organoids, we identify a collagen I-specific subset of basal-like cells that activate a mesenchymal gene program, including vimentin and ECM-degrading enzymes, while retaining epithelial junction components. We define this subset as leader cells based on four converging lines of evidence: (i) they emerge selectively in collagen I, a matrix permissive for invasion; (ii) they selectively upregulate invasion-enabling genes (e.g., *Vim*, *Flna*, ECM degradation genes); (iii) these invasion-enabling genes (e.g., *Vim*) are enriched in front-localized cells within invasive strands in organoids and patient tumors; and (iv) perturbing key effectors upregulated in these cells, namely vimentin and MMP-mediated degradation, impairs leader cell formation and collective invasion.

The selective mesenchymal program is consistent with functional properties required for basal-like cells to initiate and lead collective invasion. Vimentin, uniformly upregulated in the collagen I-specific basal subset (leader cells), has been shown to reinforce actin-rich protrusions, enhance traction force generation, and protects against mechanical stress during migration^61–63^. Vimentin positive leader cells were recapitulated in invasive carcinoma NST patient tissue, where positive basal-like cells (high K14 – low K8) at the tumor-stroma interface exhibit elongated, protrusive-like morphologies and upregulate vimentin. Despite the limitations of 2D histological analysis, these features align with the invasive front-localized phenotype observed in 3D collagen I organoids, reinforcing the relevance of basal-like leader cells in human breast tumors^64,65^. In addition to *Vim*, cytoskeletal regulators such as *Tpm2*, *Tpm4*, *Capg*, and *Flna* were enriched in the leader basal-like cell subset. These genes are known to promote actin filament stability, contractility, and mechanotransduction, collectively supporting front–rear polarity and force transmission at the leading edge^66–68^. In total, this cytoskeletal molecular repertoire may underlie directional migration through mechanically resistant environments. To overcome such barriers, our work shows that leader cells exhibit enrichment of an ECM-degradation gene signature, including *Timp1*, *Plau*, *MMP3*, *MMP13*, and *MMP14*, as well as cathepsins. This repertoire supports focal collagen I degradation, as confirmed by inhibition of both collagen proteolysis and invasion upon MMP blockade. Beyond proteolysis, leader cells express ECM-components and -modifying genes such as *Fn1*, *Tnc*, and *Loxl3*, which may contribute to broader matrix remodeling and formation of invasion-permissive paths^69–72^.

While collagen I is known to promote protrusive activity in cancer cells, its role in driving mesenchymal characteristics has remained unclear especially in collective invasion. Our data demonstrate that collagen I-induced leader cell function requires Itgα2, which mediates high-affinity adhesion to fibrillar collagen I^20^. We find that Itgα2 is essential for mesenchymal reprogramming, including vimentin expression and MMP-mediated ECM degradation. Mechanistically, Itgα2 acts as an upstream regulator of TGFβ signaling, a well-established driver of EMT^73,74^. Multiple lines of evidence support this: (i) *Inhba* is selectively upregulated in basal-like cells exposed to collagen I; (ii) *Itgα2* knockout reduces *Inhba* expression, which encodes activin A, a TGFβ ligand required for leader cell emergence; (iii) *Itgα2* loss impairs TGFβ pathway activity, as shown by reduced *Smad7* and *Pai1* expression and downregulation of mesenchymal markers such as slug and vimentin; and (iv) exogenous activin A restores TGFβ signaling and mesenchymal gene expression in *Itgα2*-deficient cells. These findings support a model in which collagen I engagement activates mesenchymal programs via Itgα2-dependent induction of *Inhba*, integrating mechanical and transcriptional signals to sustain leader cell identity.

Our data further establish activin A as a key effector of collective invasion. While supplementation with exogenous activin A promotes leader cell emergence across multiple basal-like breast cancer models, it fails to rescue invasion in *Itgα2*-deficient organoids. This suggests that although activin A can support mesenchymal traits, Itgα2 is indispensable for invasion, likely due to the dual role of α2β1 integrin in collagen adhesion and signal transduction.

Despite their mesenchymal activity, leader cells retain the expression of epithelial junction genes, including *Epcam*, *Cdh1* (E-cadherin), *Tjp2*, and *Cldn12*. An in-depth analysis of the organization and function of these junctional proteins remains to be performed. *Cdh1* upregulation observed upon *Itgα2* knockout in bulk qPCR assays must be interpreted cautiously, as these measurements reflect mixed cell populations, including luminal or non-invasive basal-like cells that may be more transcriptionally responsive to *Itgα2* loss. Nonetheless, single-cell analysis indicates that *Cdh1* transcript levels remain stable in collagen-contacting basal-like leader cells, suggesting that repressive input from *Itgα2* is either limited or buffered by concurrent epithelial-maintaining programs. This latter interpretation is further supported by the effect of TGFβ modulation: while activin A enhances mesenchymal gene expression, it does not suppress Cdh1 and instead induces a mild increase. This dual role of activin A may help explain how leader cells sustain epithelial identity despite exposure to mesenchymal-promoting cues and aligns with findings in colon^58^ and esophageal^59^ carcinoma models, where activin A supports *E-cadherin* expression. These data suggest that the collagen I-Itgα2-activin A axis enables selective mesenchymal remodeling without full junctional dissolution, reinforcing the hybrid epithelial–mesenchymal phenotype that defines basal-like leader cells during collective invasion.

The maintenance of cell-cell junction by leader cells raises the possibility that junctional integrity may contribute to mechanical coupling along the invasive strand. Cell-cell coupling may facilitate front–rear coordination and support persistent collective migration as a cohesive unit^76,78^. Thus, with such a hybrid epithelial–mesenchymal phenotype enables leader cells to remodel their environment without detachment and may confer survival advantages, including resistance to cell death and therapy, frequently observed at invasive tumor fronts^64,65^ ^80,82,84^. In contrast, single epithelial cells undergoing EMT often lose cell-cell adhesion and polarity, rendering them less effective at sustained invasion and more vulnerable to anoikis and mechanical stress^15,55^. Together, these findings define a hybrid invasion program that integrates mesenchymal dynamics with epithelial architecture, reinforcing the central role of leader cells in collective tumor progression.

The clinical relevance of this Integrin-Inhba axis is supported by our meta-analysis showing that co-expression of *INHBA*, *ITGA2*, and *ITGB1* is associated with reduced distant metastasis-free survival in grade 3 breast cancer patients. The transcriptional regulation of *INHBA* by Itgα2 remains to be fully defined, but may involve YAP, a mechanoresponsive transcriptional co-activator active in leader cells and previously shown to control *INHBA* expression^75^.Beyond Itgα2 and activin A, other integrin-dependent mechanisms likely contribute to ECM-mediated TGFβ activation and EMT^77,79^. Latent TGFβ ligands stored in the ECM require mechanical activation, a process mediated by integrins such as αvβ6 and αvβ8 that apply tension to the latency-associated peptide (LAP) to release active TGFβ^77^. Traction forces generated by leader cells may further promote this process, linking ECM remodeling to the reinforcement of mesenchymal traits. Notably, while ECM degradation is independent of TGFβ signaling, it remains dependent on Itgα2, suggesting alternative regulatory axes. Whether Itgα2 modulates MMP activity at the transcriptional or post-transcriptional level remains to be determined^81^.

Our findings demonstrate that basal-like leader cells inhabit a transcriptionally defined niche in which ECM engagement and autocrine/paracrine signaling converge to support mesenchymal traits and collective invasion. Cytoskeletal remodeling, ECM degradation, and retention of epithelial junction gene expression appear to function in parallel to maintain leader cell identity. This coordination enables basal-like cells to simultaneously interact with and remodel their environment while maintaining cohesion within the invading strand. Although the precise regulatory interactions between these modules remain to be defined, the data support a model in which collagen I-Itgα2 signaling reinforces the partial EMT state required for collective invasion. Targeting this axis may represent a therapeutic strategy to disrupt leader cell–mediated invasion in aggressive epithelial cancers.

## Materials & Methods

### Antibodies and reagents

The following primary antibodies were used: rabbit anti-human vimentin (Cat#ab92547, Abcam), chicken anti-human vimentin (Cat#PA1-16759, Invitrogen) rabbit anti-human keratin 14 (Cat# 905301, Biolegend), rat anti-mouse keratin 8 (Cat# 531826, DSHB), rabbit anti-rat collagen I cleavage site (Col ¾, Cat#0217-025, immunoGlobe), rabbit anti-human integrin α2 (Cat#ab181548, Abcam), rabbit anti-human Inhba (Cat#10651-1-AP, Proteintech), mouse IgG1 isotype control (MAB002, R&D Systems), anti-activin A antibody (Cat#AF338, R&D Systems). The following secondary antibodies were used: Alexa fluor-488/568/647-conjugated goat anti-mouse, -rat, or -rabbit antibody (Invitrogen). To visualize the nucleus and F-actin, 4ʹ,6-diamidino-2-phenylindole (DAPI, Cat#D9564, Sigma) and Alexa-fluor-568-conjugated phalloidin (ThermoFisher) were used, respectively.

The following reagents were used: Ilomastat/GM6001 (Cat#2983, TOCRIS), SB431542 (Cat#616461, Merck), A83-01 (Cat#2939/10, Bio-Techne), activin A (Cat#AF338, R&D Systems), BSA (Cat#A9647, MERCK), and Dimethyl Sulfoxide (Cat# 5.89569, MERCK).

### Organoid culture

The MMTV-PyMT and *Apc*^1572T^ organoid lines were generous gifts from Jacco van Rheenen^83^ and R.F.^33^, respectively. The 4T1 metastatic breast cancer cell line was used (CRL-2539 ATCC). All cells (organoids and 4T1) were cultured in 20-25μl drops of cultrex reduced growth Factor basement membrane extract (BME) (Cat# 3533-005-02, RGF type 2 Path Clear, Bio-Techne) as previously described^85^. Similar to the organoids (MMTV-PyMT, *Apc*^1572T^), the 4T1 cells grew in BME into rounded multicellular structures (spheroids). The growth medium of MMTV-PyMT organoid line and 4T1 spheroids was composed of dulbecco’s modified eagle medium F12 (Cat# 12634010, Gibco), supplemented with 10uM HEPES (Cat#15630080, Gibco), 10 U/ml Pen-Strep (Cat#17-602E, Lonza), 1% GlutaMAX 100X (Cat# 35050061, Life Technologies), 125uM N-Acetyl-L-Cysteine (Cat#A9165, Sigma), 0.01% Primocin (Cat#ant-pm-2, Bio-Connect), 1x B27 (Cat#17504044, Life Technologies), and 2.5 nM human FGF2 (42,5 ng/ml)(Cat# 100-18B, PeproTech). *Apc*^1572T^ organoids were grown in dulbecco’s modified eagle medium F12 supplemented with 10 U/ml Pen-Strep, 1x B27, 0.01% Primocin, 1.16 nM FGF2 (20ng/ml), and 3.2nM EGF (20ng/ml, Cat#AF-100-15, PeproTech).

Cultures were maintained at 37°C, 5% CO2, and humidified atmosphere. They were passaged approximately twice a week by dissociating into single cells using trypsin-EDTA (TrypLE Express, Cat# 12605-010) for a maximum duration of 5 min at 37°C. The organoids and spheroids were further mechanically dissociated into single cells by repeated pipetting. The single cells were washed with DMEM/F12 (+HEPES, GlutaMAX and pen-strep) followed by centrifugation (260G, 5min. 4°C). Cell pellets were suspended in BME. After polymerization, the gels were supplemented with culture media.

### Collagen I invasion assays

Confluent BME cultures containing medium-sized breast cancer organoids (MMTV-PyMT and *Apc*^1572T^) and spheroids (4T1) were collected by resuspending with cold media (DMEM/F12 supplemented with HEPES, Pen-Strep, and 1% GlutaMax). Organoids were spun down for 4 minutes at 260 G (4°C), and supernatant was discarded. To remove remaining BME, the pellet was incubated with dispase 1mg/ml (Cat#17105-041, Life Technology) for 15-minute incubation at 37°C water bath. This was followed with 4 washing rounds. Next, the organoids were embedded in 40 μl drops of collagen I (2 mg/ml) or BME gels in 24 or 48 -well plates. Non-pepsinized rat-tail collagen (Cat#354236, Corning) was prepared at a final concentration of 2 mg/ml using 10X phosphate buffer saline (PBS) (Gibco), NaOH (final pH of 7.5) and dH2O according to the manufacturer’s instructions. Collagen I was pre-polymerized for 2 hours on ice, then organoids were added to the collagen mix followed by 5 min. incubation at room temperature and plating on pre-warmed plates (Costar) and incubated for 30 minutes at 37°C to polymerize^85^. Subsequently, medium was added.

### Single cell sequencing analysis

Previously published data from GEO (GSE197821)^27^ was analyzed in R using Seurat V4^86^. Following data integration, cells were clustered into 5 subgroups using shared nearest neighbor (SNN) with resolution parameter 0.3. Gene ontology analysis was performed using the EnrichR package^87^ based on the hallmark gene sets from the molecular signature database^88^. For this analysis, differentially expressed genes were identified by comparing the distinct basal populations with FindMarkers, and filtering genes using cutoffs for pvalue (<0.01) and log2FC (>0.5). Other signatures (Supplementary information 1) were evaluated using the AddModuleScore function, and visualized with a violin plot across the distinct cell populations or log2FC values for a two-group comparison were directly visualized as bar plots.

### Brightfield microscopy

The breast cancer organoids and spheroids were imaged using the EVOS M5000 microscope using the 2x and 10x objective (2x objective Numerical Aperture (NA) = 0.08, 10x objective, NA=0.4). For invasion quantification, a region of interest (ROI) was manually drawn around the organoid body, and invasive strands were traced by drawing a line from the organoid border until the tip of the strand using ImageJ (ImageJ; 1.40v; National Institutes of Health). The organoid body area (in µm²), number of pointed tips (leader cells) and strand length (in µm) were measured^55^.

### Immunofluorescence and confocal microscopy

Organoids and spheroids cultured in 3D collagen I were fixed using 4% paraformaldehyde (Sigma-Aldrich) (15-minute incubation at room temperature (RT)), followed by three washing rounds using 1X PBS (Cat#D8537, Sigma-Aldrich). After washing, the collected gels were transferred into 1.5 mL Eppendorf tubes and blocked using 10% normal goat serum (Gibco) with 0.3% Triton-X (Sigma-Aldrich) in 1X PBS and incubated for 1 hour while shaking at RT. This was followed with primary antibody incubation in 0.1% bovine serum albumin (BSA; Sigma Life Sciences) with 0.3% Triton in 1X PBS. Samples were incubated with the primary solution overnight at 4 °C with shaking. Dilutions of primary antibodies used in this study include: K14 antibody (1:400), K8 antibody (1:50), vimentin antibody (1:200), collagen ¾ antibody (5 µg/mL), Itgα2 antibody (1:400), Inhba antibody (1:70). After removing the primary antibody solution, the samples were washed four times 15 minutes in 1X PBS. After washing, the samples were incubated with the secondary antibody solution containing Alexa-fluor 488/568/647 conjugated secondary antibodies together with DAPI (5µg/ml) and Phalloidin (1:500) overnight at 4 °C with shaking. The next day, the gels were thoroughly washed by four cycles of 10 minutes 1X PBS. For collagen ¾, similar staining steps were performed but the blocking and antibody buffers did not contain Triton. The samples were imaged on glass bottom WillCo Wells (WillCo Well BV) using confocal microscopy (LSM 880 Zeiss) with a 40X magnification objective (NA=1.1).

To quantify vimentin expression in K14⁺ cells (basal-like cells), nuclear (DAPI) and cytosolic (F-actin and keratin 8) segmentation was performed using Cellpose^89^. Mean gray values for vimentin were measured in nuclear and cytosolic compartments, and after background subtraction, cytosolic expression was quantified by subtracting the nuclear signal from the cytosolic signal; negative values were set to zero. Leader and rim cells were manually annotated based on spatial criteria: leader cells were located at the tip of an invading strand; lateral leader cells were located within the invasive strand and in contact with the ECM; and rim cells were located at the outer layer of the organoid body in contact with the ECM.

To analyze the effect of *Itgα2* KO on Inhba expression, a single confocal z-slice from the mid-plane of WT and KO organoids was used. Using Fiji, contours were manually drawn around the cytosol and nucleus (based on Phalloidin and DAPI signals) of basal-like cells (low K8) that are located at the organoid-ECM interface. The mean gray value of Inhba intensity was measured on 8-bit images, with background correction performed by subtracting the intensities of cell-free regions. The collagen ¾ signal quantification was performed using an ImageJ macro that starts by selecting the middle z-layer from the acquired confocal stack. The script then applies a threshold (adjusted manually) to the F-actin signal to create a binary mask, selecting the organoid while excluding the surrounding ECM and background. Using this mask, the macro generates an outline (line width = 15) forming the border of the organoid and defining the region of interest. Collagen ¾ and collagen I reflection signal are then measured within the outlined regions and the relative ratio of collagen ¾ to collagen I reflection signal is calculated.

### Immunofluorescent staining of patient invasive carcinoma NST samples

Immunofluorescence was performed on 8 µm whole invasive breast cancer tissue sections diagnosed as high-grade invasive carcinoma NST (Table 3). The use of material was approved by the Tissue Science Committee of the University Medical Center Utrecht and informed consent was obtained from the participants. Tissues were deparaffinized with Xylene (2X, 2 minutes) followed by hydrating the tissues using a sequential descending percentage of ethanol (100%, 96%, 70%, and 50%; 2X, 2 minutes each). Antigen retrieval was performed using Tris-EDTA buffer (15 min, 95–100 °C) followed by immunostaining as previously described^55^. Briefly, tissue sections were blocked using 5% normal goat serum in 0.05% Tris-buffered saline (TBS) with Tween™ 20 detergent (TBST) for 1 h at RT followed by primary antibody incubation at 4 °C (overnight) using the following antibody dilutions: Itgα2 (1:300), vimentin (1:300), and K8 (1:50). After antibody incubation, samples were washed four times (TBST, 5 min each) and incubated with secondary Alexa Fluor 488/546/647-conjugated goat anti-mouse, anti-rabbit, anti-rat antibodies and with DAPI (1 μg/mL), for 1 h at room temperature. Samples were washed four times (TBST, 5 min each) and mounted with Immu-Mount (Epredia). Imaging was performed using confocal microscopy (LSM 880 Zeiss) with a 40x objective (NA =1.1). For the analysis of Itgα2, K8, K14, and vimentin expression in tumor cells, cancer cell groups were first identified based on their K8-positive cells organized into protrusive multicellular structures. Cells were classified as luminal-like (K8-high), basal-like (K14-high), or stromal (spindle-shaped with elongated nuclei). Using Fiji, contours were manually drawn around the cytosols of selected cells using a merged channel combining all cytoskeletal markers (K8, K14, and vimentin). The mean gray value of cytoskeletal markers and Itgα2 intensity was then measured within the selected regions of interest and background-corrected by subtracting the intensity of a cell-free region.

### Generation of doxycycline inducible knockdown organoid models

To knockdown vimentin, MMTV-PyMT and *Apc*^1572T^ organoids expressing doxycycline-inducible shRNA against vimentin were generated. Two independent shRNA sequences were designed (Table 1). Successful incorporation of the shRNA sequences was confirmed by agarose gel electrophoresis, showing digestion products, and by DNA sequencing. Lentiviral particles were then produced by transfecting HEK293 cells with the FUTG-tetInd-shRNA vector and packaging plasmids using X-tremeGENE (Cat#6365809001, Sigma Aldrich).

**Table 1.**
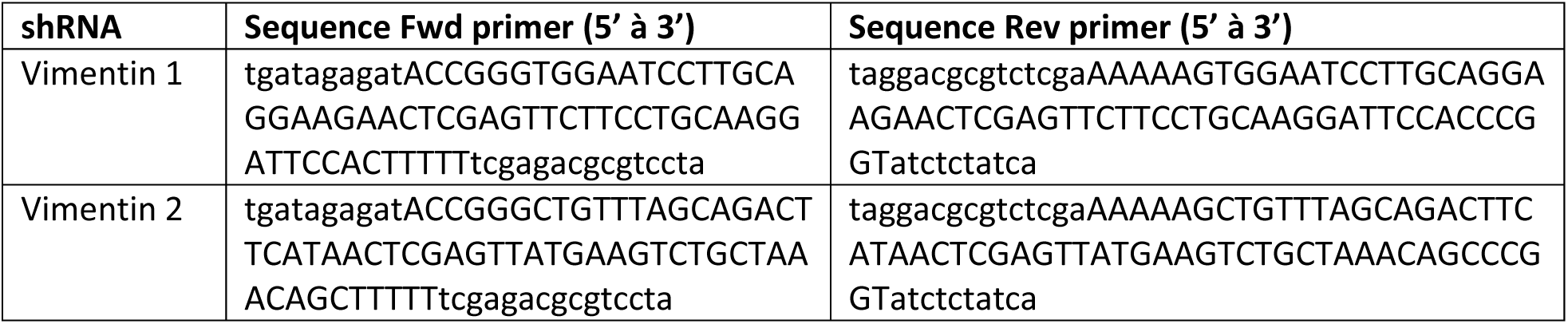
Sequences of shRNA constructs (1 and 2) targeting vimentin.

Supernatant containing viral particles from HEK293 cell cultures were collected at two time points, 48 and 72 hours following transfection. To concentrate the viral particles, the supernatant was spun down by ultra-centrifugation (175000G for 2.5 hours), using the Optima LE-80K Ultracentrifuge. For organoid infection, organoids were first dissociated into small clusters of cells and single cells, then incubated with concentrated lentiviral particles for 4–6 hours. Following transduction, cells were re-embedded into BME and maintained for 48 hours. Transduced organoids were enriched by adding the selection marker blasticidin (10 µg/mL) (Cat# ANT-BL-05, Invitrogen). To induce vimentin knockdown, doxycycline (2 µg/mL, Cat# D9891, Sigma-Aldrich) was added to the organoid cultures in BME for 3 days. After pretreatment, organoids were extracted from BME and seeded into collagen I matrices to assess for effects on invasion and knockdown efficiency (Western blot analysis and/or immunofluorescence microscopy).

### Generation of *Itgα2*-deficient organoids

Itgα2 knockout MMTV-PyMT organoids were generated by transfection with a pX458 vector that expresses Cas9-T2A-GFP and Itgα2 sgRNAs targeting the first coding exon of Itgα2 (sgRNA1 gAATTGTCTGGCGTATAATGT, sgRNA2 cGCGTATAATGTTGGCCTCCC). To transfect the organoids, 6 µg of the pX458 vector was mixed with 300 µL Optimem and 18 µL TransIT transfection reagent (Cat# MIR 2300, Mirus) and incubated for 15 minutes at room temperature to allow complex formation. The transfection mixture was then added to single-cell suspension in culture medium (5% FCS) and plated on 3 wells of a 6-well plate for 24 hours. Next, the culture medium was replaced with the FCS-free organoid culture medium. The following day, GFP-positive cells were sorted using a FACS Aria II flow cytometer and were embedded in BME and maintained in culture until the single cells developed into multicellular organoids. Around 10 single organoids were picked from the BME gel using a pipette under a brightfield microscope. These selected organoids were trypsinized and expanded separately in BME gels to establish monoclonal lines. To confirm Itgα2 knockout, western blot analysis DNA sequencing were performed. For sequencing, genomic DNA was isolated from organoid pellets using the QIAamp DNA Micro Kit (Cat# 56304, Qiagen). A 999 bp region around the Cas9 target site was amplified in a 50 µL reaction containing 0.5 µLQ5 Polymerase (Cat# NEB M0491L, Bioke), 10 µL 5x Q5 Buffer (Bioke), 1 µL isolated gDNA, 500 nM forward primer Itgα2: TCCAGAGCCTTCCCTTACAA, 500 nM reverse primer Itgα2: AGCCCTGCAGAAAGACGATA, 10 µL Q5 high GC buffer (Bioke), 200 µLM dNTP’s (Roche) and 17.5 µL H2O. The PCR program was as follows: initial denaturation at 96°C for 2 min, denaturing at 95°C for 30 seconds, annealing at 57.9°C for 30 seconds extension at 72°C for 45 seconds, and final extension at 72°C for 7 minutes. The denaturing, annealing, and extension steps were repeated 32 times. The PCR products at around 999 bp were isolated from TAE agarose gels using the QIAquick Gel Extraction Kit (Cat# 28704, Qiagen) and sent for sequencing (Macrogen) using the forward primer.

### Western blot analysis

To isolate organoids for whole cell lysates, collagen I gels were first digested with collagenase mixture (0.56 mg/mL; Cat# C0130, Sigma-Aldrich) for 3 minutes at 37°C with gentle shaking^85^. Following digestion, organoids were washed with PBS to remove residual collagenase.

After washing with PBS, organoids were lysed in SDS sample buffer (62.5 mM Tris HCl, 2% SDS, 10% glycerol, 50 mM DTT and bromophenol blue). Lysates were sonicated for 5 cycles at 4°C (Bioruptor) followed by 5-minute boiling at 95°C. Samples were loaded onto a 10% acrylamide gel and separated by SDS-PAGE in Tris-glycine running buffer at 100V for ∼2 hours. Proteins were then transferred onto a Polyvinylidene difluoride (PVDF) membrane (Cat# IPFL00010, Millipore) using wet transfer at 0.35 A for 90 minutes. To reduce non-specific antibody binding, membranes were blocked for 1 hour at RT in 5% non-fat dry milk dissolved in 1X TBS. Blots were incubated overnight at 4°C with primary antibodies in antibody solution containing 0.1% Tween-20 and 5% BSA in TBS. After extensive washing (4X 10 minutes, 0.1% Tween-20 in TBS), membranes were incubated with secondary antibodies for 1 hour at RT followed by a second round of washing. The following dilutions of primary and secondary antibodies were used Itgα2 (Cat#ab181548, Abcam; 1:5000), vimentin (Cat#ab92547, Abcam; 1:2500) and glyceraldehyde-3-phosphate dehydrogenase (GAPDH) (Cat#MAB374; 1:5000). Secondary antibodies used were Odyssey goat anti-mouse 800 and goat anti-rabbit 680 (1:2500). Fluorescent signal was measured and visualized using the Amersham Typhoon scanner. To calculate knockdown efficiency using western blot, mean gray values were measured using FIJI software for both the protein of interest and the housekeeping gene (GAPDH). The efficiency was determined using the following formula:

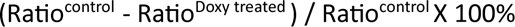

The ratio was calculated by dividing the mean grey values of the protein of interest (band) by the corresponding mean gray value of the GAPDH band.

### Real-Time PCR (qPCR)

For RNA isolation from organoids embedded in collagen I or BME, gels were collected and lysed by mechanical disruption in 500 μL TRIzol™ reagent (Thermo Fisher) per 40 µL gel. RNA was extracted using the NucleoSpin® RNA kit (Cat#740955.50, Macherey-Nagel) and subsequently converted into cDNA using the iScript™ cDNA Synthesis Kit (Cat#1708891, Bio-Rad) (500-1000 ng in 20 µL).

For qPCR, the reaction mixture contained cDNA, 0.4 µM forward and reverse primers, and 1X FastStart Universal SYBR Green Master mix (Cat#4913850001, Roche) to a final volume of 15 µL.

qPCR primers (Table 2) were designed to generate products between 50-150 bp and were confirmed to specifically bind the mouse gene of interest using the NCBI n-BLAST tool. Primers were validated through cDNA titration and were approved when amplification efficiency ranged between 80-120% and when a single distinct melting peak was observed during the qPCR melting curve analysis

**Table 2.**
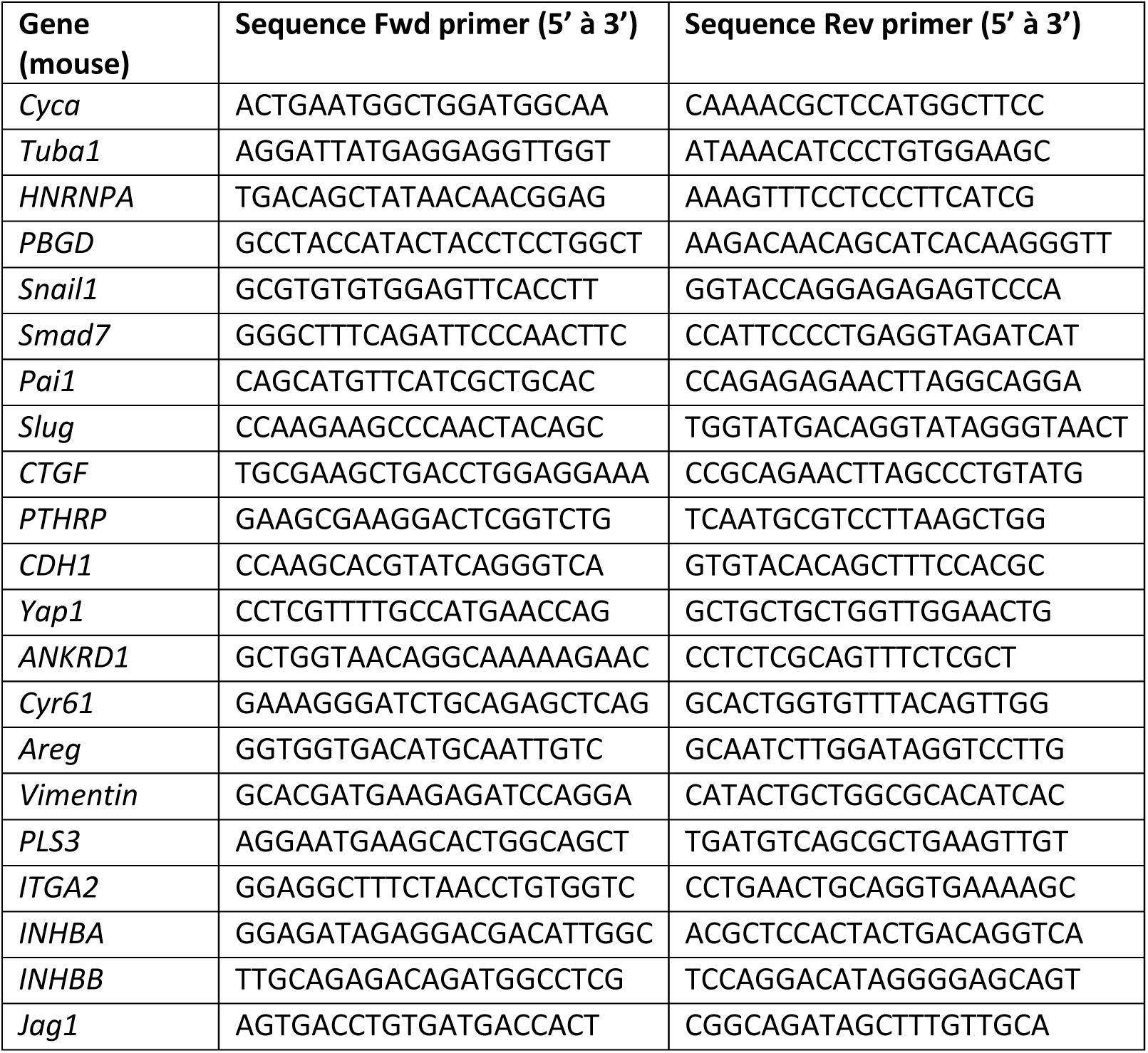
Sequences of qPCR primers.

**Table 3.** Patients’ information.

| **Patients** | **Diagnosis** | **Subtype** | **Age** | **Grade** |
| --- | --- | --- | --- | --- |
| 14 | Invasive carcinoma NST | ER (-) / PR (-) / HER2 (+) | 74 | 3 |
| 25 | Invasive carcinoma NST | ER (+) / PR (+) / HER2 (-) | 47 | 2 |
| 19 | Invasive carcinoma NST | ER (+) / PR (+) / HER2 (-) | 37 | 3 |
| 45 | Invasive carcinoma NST | ND | 42 | 3 |
| 5 | Invasive carcinoma NST | ER (+) / PR (-) / HER2 (-) | 39 | 3 |

Age: age at primary diagnosis
NST: no special type
ND: Not Determined

The qPCR was performed using a thermal cycler (CFX384, Bio-Rad) using the following program: 95 °C for 10 min, followed by a 40x repeated cycle of 95 °C for 10 s (denaturation), 55 °C for 10 s (annealing) and 72 °C for 30 s (extension). A melt curve analysis was conducted with a program of 95 °C for 10 s, followed by incremental increases of 0.5 °C from 65 °C to 95 °C to confirm primer specificity and detect potential non-specific amplification. mRNA expression levels were normalized across samples using the average expression of four housekeeping genes (*Cyca*, *Tuba1*, *Hnrnpa*, and *Pbgd*). Data was analyzed using the CFX Maestro software, and relative gene expression changes were calculated using the ΔΔCt method.

### Statistical analysis

For statistical analysis, Graph Prism (version 8.0) was used. The two-tailed unpaired Mann–Whitney (two groups) or Kruskal–Wallis test (more than two groups) with Dunns’ correction were used for statistical analysis. For qPCR analysis, parametric statistical analysis was used after confirming normal distribution of the data by Shapiro-Wilk test. P values less than 0.05 were considered significant.

## Acknowledgments

This work was supported by grants from the Dutch Cancer Society (KWF 2020-13552, AAK), the European Union’s Horizon 2020 FET Proactive program (grant agreement No. 731957, MECHANO-CONTROL, AAK), and ZonMw (Vici 9150181910029). We thank Kitty van Zwieten, Lars Kemp, Marjolein Vliem, Livio Kleij, Mirjam van der Net, and Ingrid Verlaan for the technical support, and Dr. Johan de Rooij, Dr. Fried Zwartkruis, Dr. Eric Kalkhoven, and Dr. Paul Coffer for scientific discussions.

## Author contributions

A.G.E.G., L.J., and A.A.K. designed the study. A.G.E.G., L.J., H.K., S.S., and M.M. performed the experiments. M.P.V. performed bioinformatics analysis. A.G.E.G., L.J., S.S., H.K., M.M., I.dS, and A.A.K. analyzed the data. L.J., D.W., H.K., R.F. established and provided the organoid lines. A.G.E.G. and A.A.K. wrote the manuscript. All authors read the manuscript and were given the opportunity to provide input.

## Supplementary Figures

**Supplementary Figure 1.**
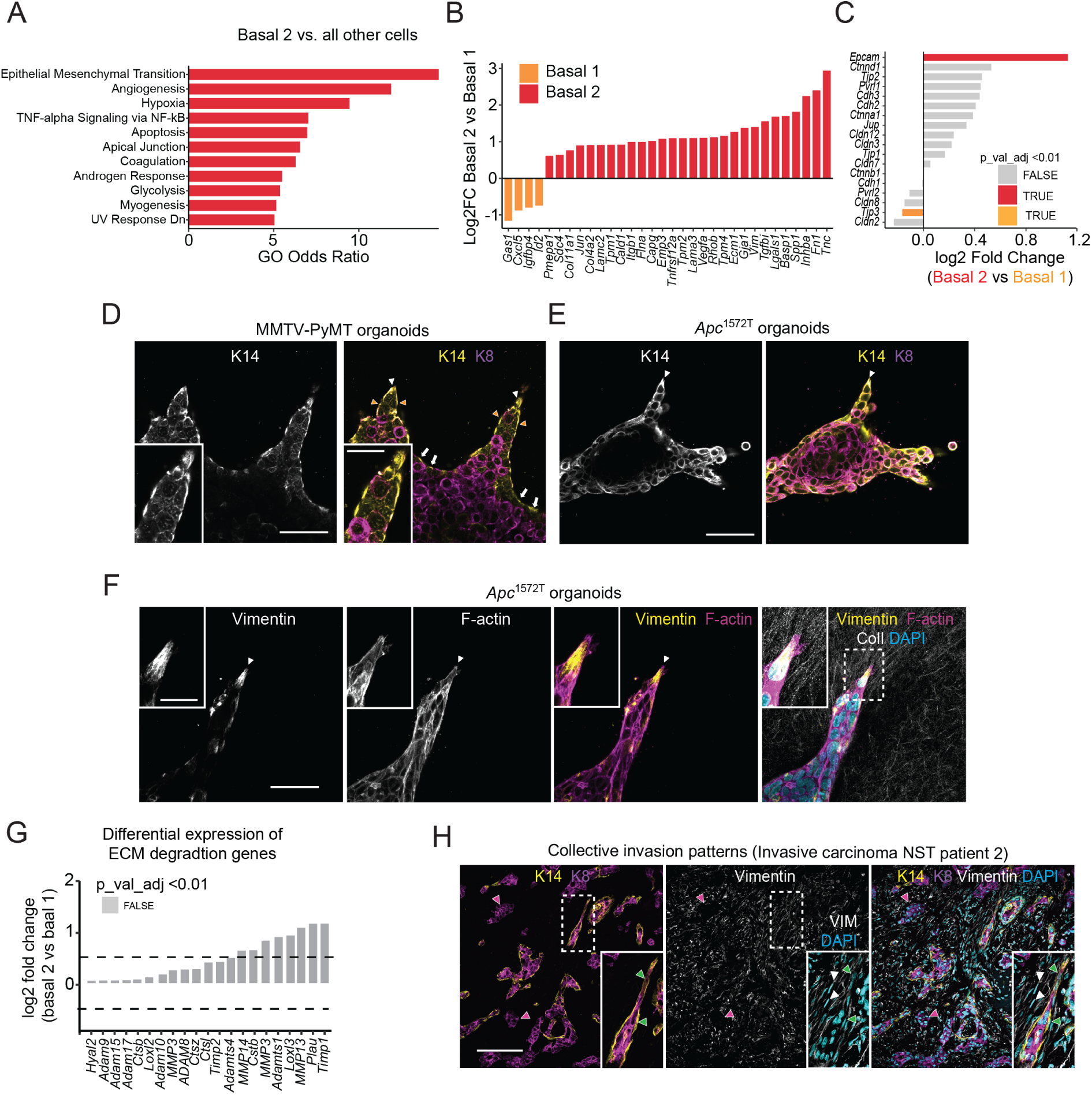
Basal-like leader cells exhibit mesenchymal traits while maintaining epithelial junctions during collective invasion. **(A)** Gene set enrichment analysis (GSEA) of basal 2 clusters showing significantly enriched pathways compared to all other cells (filtered for NES > 0.5 and P < 0.01). **(B, C)** Log₂ fold changes in gene expression between basal 2 and basal 1 subsets, based on scRNA-seq data from MMTV-PyMT organoids**. (B)** Top 28 differentially expressed genes. **(C)** Genes within a curated list of known cell–cell junction components, including adherens, tight, gap, and desmosomal junction proteins. **(D, E)** Confocal imaging of K14 and K8 in **(D)** MMTV-PyMT and **(E)** *Apc*^1572T^ organoids embedded in 3D collagen I for 3 days. White arrowheads depict protrusive basal-like leader cells (K14-high, K8-low). **(F)** Vimentin immunostaining in *Apc*^1572T^ organoids embedded in 3D collagen I for 3 days, arrowheads depict vimentin in protrusive leader cells. **(G)** Log₂ fold changes in the expression of ECM-degrading and remodeling genes between basal 2 and basal 1 cells from scRNA-seq of MMTV-PyMT organoids. The ECM degradation gene set comprises matrix-modifying and proteolytic enzymes, including matrix metalloproteinases (MMPs), ADAM and ADAMTS family members, cathepsins, serine proteases, hyaluronidases, lysyl oxidases, and associated regulators (e.g., TIMPs, serpins). None of the observed differences reached statistical significance (adjusted P ≥ 0.01). **(H)** Immunostaining of K14, K8, and vimentin in a whole tissue section from a K14-positive invasive carcinoma NST patient, with zoom-in on protrusive cell co-expressing K14 and vimentin. Arrowheads: green, basal-like cells; magenta, luminal cells; white, fibroblast-like cells (elongated, spindle-shaped). Scale bars: 50 µm **(D, E, F)**; 10 µm (**D**, **F**, zoom-in); 100 µm **(H)**; 25 µm (**H**, zoom-in).

**Supplementary Figure 2.**
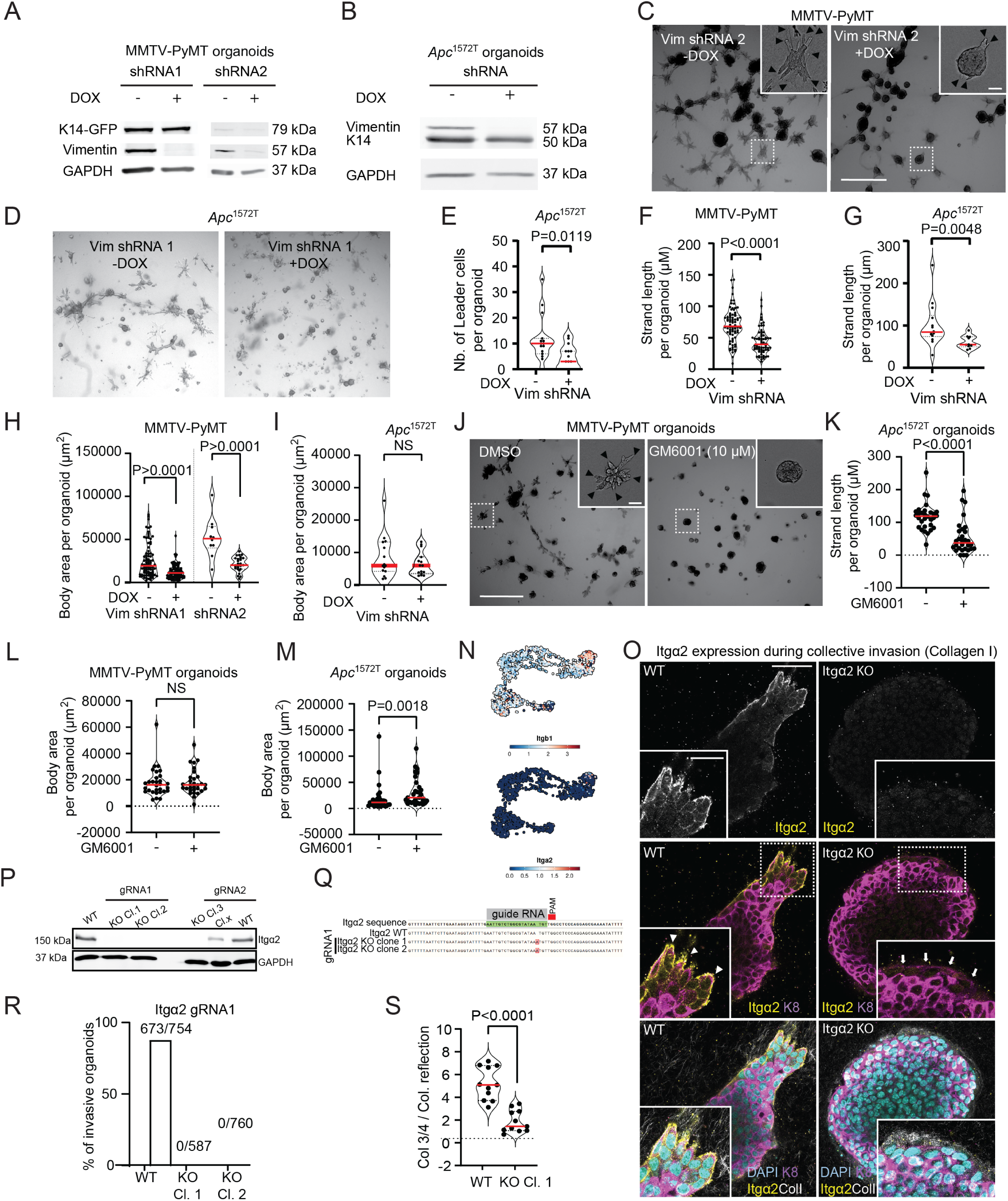
Vimentin and integrin α2 are required for leader cell emergence and collective invasion in breast cancer organoids. (A, **B)**Western blot analysis of vimentin, K14, and GAPDH from whole-cell lysates of **(A)** MMTV-PyMT and (B) *Apc*^1572T^ organoids expressing doxycycline-inducible vimentin shRNAs cultured in collagen I for three days with or without doxycycline (DOX). **(C, D)** Brightfield imaging of control and organoids with vimentin knockdown (day 3, collagen I). Arrowheads **(C)**: leader cells. **(E-I)** Quantification of **(E)** number of leader cells per organoid, **(F, G)** average length of invasive strands, and **(H, I)** organoid surface area in control and vimentin knockdown organoids: **(F, H)** MMTV-PyMT: *n* = 87 (–DOX), *n* = 87 (+DOX) for shRNA1, and *n* = 29 (−DOX), *n* = 44 (+DOX) for shRNA2, from three (shRNA1) and two (shRNA2) independent experiments. (E, G, I) *Apc*^1572T^: *n* = 14 (−DOX), *n* = 10 (+DOX) organoids (shRNA1) from one experiment. **(J)** Representative brightfield images of MMTV-PyMT organoids in collagen I treated with GM6001 or DMSO control for three days. Arrowheads: invasive strands. **(K-M)** Quantification of **(K)** average length of invasive strands and **(L, M)** organoid surface area in organoids treated with GM6001 or DMSO control for three days: (K, M) *Apc*^1572T^ and **(L)** MMTV-PyMT organoids: *n* = 30 organoids per condition from three independent experiments. **(N)** UMAP plots of *Itgβ1* and *Itgα2* mRNA expression across the epithelial subpopulations. **(O)** Confocal imaging of Itgα2 and K8 in MMTV-PyMT organoids with Itgα2 WT or knockout (KO) grown in collagen I for 3 days. White arrowheads: invading strands in Itgα2 WT organoids led by basal-like cells (low K8, high Itgα2, insets), White arrows: non-invading basal-like cells at the ECM interface in Itgα2 KO organoids (low K8, low Itgα2, insets). **(P)** Western blot analysis showing Itgα2 and GAPDH expression from whole cell lysates of MMTV-PyMT organoids WT or KO (gRNA1 or gRNA2). **(Q)** DNA sequence of the *Itgα2* gene to confirm gene knockout by Crispr-Cas9 gene editing. Red regions indicate the insertion of one base pair compared to the wildtype *Itgα2* sequence. **(R)** Percentage of organoids exhibiting one or more invasive strands in Itgα2-WT and Itgα2-KO (clones 1 and 2 from gRNA1) MMTV-PyMT organoids. **(S)** Mean gray value of col ¾ relative to collagen I reflection in MMTV-PyMT organoids with Itgα2-WT and Itgα2-KO. Red line indicates the median, from *n* = 11 organoids per group from two independent experiments. Scale bars: 1000 µm **(C, D, J)**, 10 µm (**C**, **J**, zoom-in), 50 μm **(O)**, 25 μm (**O**, Zoom-in). P values: two-sided unpaired Mann–Whitney test **(E-I, K, L, M, S)**.

**Supplementary Figure 3.**
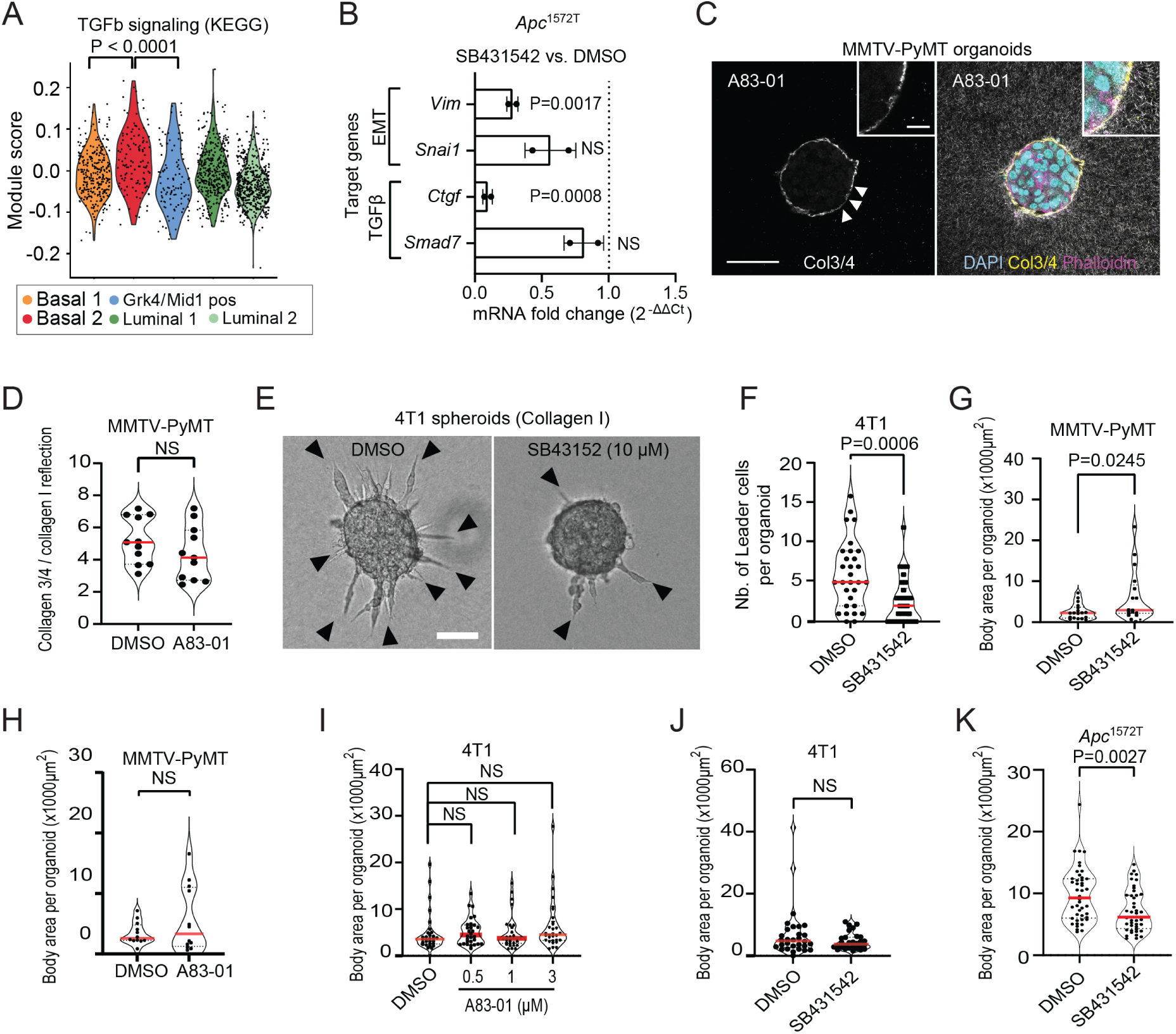
TGFβ signaling promotes mesenchymal traits and leader cell emergence during collective invasion. **(A)** Module scores representing active TGFβ signaling based on the KEGG gene set reflecting relative pathway enrichment in basal 2 compared to the other subpopulations identified by single-cell RNA sequencing. **(B)** qPCR analysis of classical TGFβ and EMT target gene expression in *Apc*^1572T^ organoids cultured in 3D collagen I for three days. Values represent mean normalized mRNA expression (relative to housekeeping genes) in organoids treated with 5 μM SB431542, shown relative to DMSO-treated controls (dashed line). Data represent mean ± SD from two independent experiments. **(C)** Confocal imaging of col ¾ and F-actin in MMTV-PyMT organoids in 3D collagen I treated with A83-01 (0.5 μM) for 1 day. Arrowheads: pericellular signal of col ¾ (degraded collagen I). Inset, organoid-collagen I interface. **(D)** Mean gray value of col ¾ relative to collagen I reflection in MMTV-PyMT organoids after treatment with A83-01 or DMSO control. Red line indicates the median, from *n* = 11 organoids per group from two independent experiments. **(E)** Brightfield images of 4T1 spheroids cultured in collagen I for two days, treated with 10 μM SB431542 or DMSO control. Black arrowheads: leader cells. **(F)** Number of leader cells per spheroids in treated (10 μM SB431542) and control conditions (DMSO). Red line indicates the median, from *n* = 30 organoids per condition from three independent experiments. **(G-K)** Surface area of organoid body after treatment with SB431542 or DMSO control in **(G)** MMTV-PyMT organoids, **(J)** 4T1 spheroids, and **(K)** *Apc*^1572T^ organoids, or treatment with A83-01 or DMSO control in **(H)** MMTV-PyMT organoids and **(I)** 4T1 spheroids by a serial dilution (0.1 μM, 1 μM, and 3 μM). Red line indicates the median from **(E)** *n =* 18 organoids, **(F)** *n =* 12 organoids, **(G)** *n =* 44 organoids, **(H)** *n =* 30 spheroids from three independent experiments. Scale bars: 50 μm **(C)**; inserts 10 μm (**C**, Zoom in); 100 μm **(E)**. P values, one-way ANOVA with Tukey’s test for multiple comparisons **(A)**; two-sided Student’s *t* test **(B)**; two-sided unpaired Mann–Whitney test **(D, F, G, H, J, K)**; two-sided Kruskal-Wallis test (Dunn’s multiple comparison) **(I)**.

**Supplementary Figure 4.**
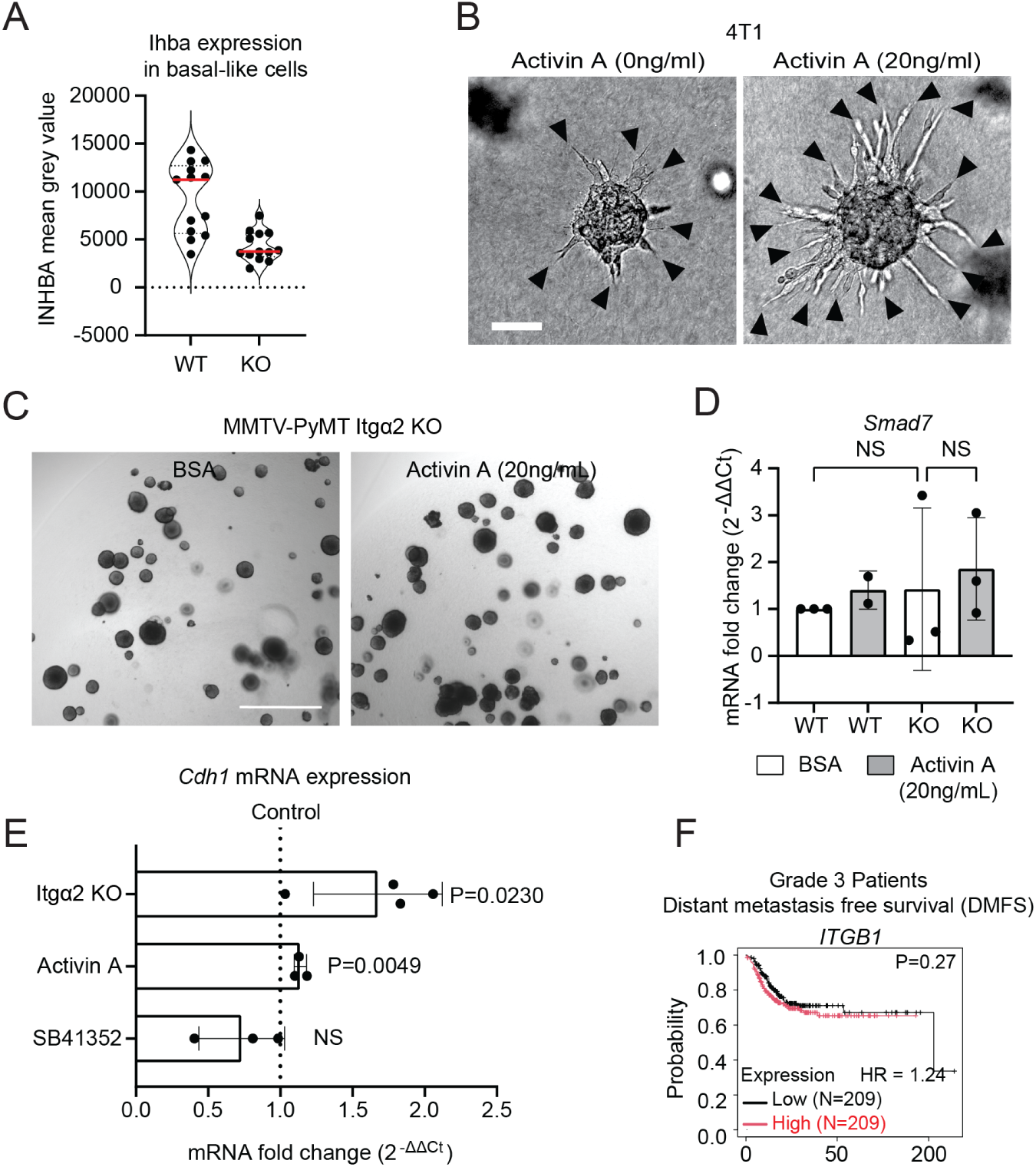
Integrin α2 regulates *INHBA* expression and TGFβ pathway activation to support leader cell function and is associated with poor prognosis in breast cancer. **(A)** Mean-gray value of Inhba in basal-like cells (K8-low) located at the rim of Itgα2-WT vs. Itgα2-KO MMTV-PyMT organoids (3D collagen I, day 1). Red line indicates the median, from *n* =11 cells from 6 Itgα2-WT organoids, and *n* = 12 cells from 7 Itgα2-KO organoids from one experiment. **(B, C)** Brightfield images of **(B)** 4T1 spheroids and **(C)** MMTV-PyMT organoids (Itgα2-WT or Itgα2-KO) cultured in collagen I for **(B)** two or **(C)** three days, treated with activin A ligand (20ng/μl) or an equal volume of 0.1% BSA in PBS as control. Black arrowheads: invasive strands **(B)**. **(D)** qPCR analysis of *Smad7* mRNA expression in Itgα2-KO versus Itgα2-WT MMTV-PyMT organoids treated with activin A (20 ng/μl) or vehicle control (0.1% BSA) for three days. Bar graphs represent mean normalized expression values ± SD from three to four independent experiments. **(E)** PCR analysis of *Cdh1* mRNA expression in Itgα2-KO relative to Itgα2-WT, activin A-treated relative to BSA control-treated, and SB41352-treated relative DMSO-control in MMTV-PyMT organoids cultured in 3D collagen I for three days. Values represent mean normalized mRNA expression (relative to housekeeping genes). Data are presented as mean ± SD from three to four independent experiments. **(F)** Kaplan–Meier analysis correlating high vs. low mRNA expression of *ITGB1* with distant metastasis-free survival (DMFS) in patients with grade 3 breast cancer. Scale Bars: 100 μm **(B)**, 1000 μm **(C)**. P values, one-way ANOVA with Šidák’s test for multiple comparisons **(D)**; two-sided Student’s *t* test **(E)**; Log-rank test **(F)**.

**Supplementary Movie 1.** Time-lapse confocal imaging of MMTV-PyMT organoids expressing endogenously tagged K14-GFP and Histone 2b (H2b)-mScarlet showing initiation and elongation of collective invasion in 3D collagen I (reflection).

Field size of a single confocal slice: 354 μm x 354 μm; time interval = 1 hour.

